# African swine fever virus-like integrated elements in a soft tick genome – an ancient virus vector arms race?

**DOI:** 10.1101/2020.03.08.978106

**Authors:** Jan H. Forth, Leonie F. Forth, Samantha Lycett, Lesley Bell-Sakyi, Günther M. Keil, Sandra Blome, Sébastien Calvignac-Spencer, Antje Wissgott, Johannes Krause, Dirk Höper, Helge Kampen, Martin Beer

## Abstract

**Background:** African swine fever virus (ASFV) is the only known DNA-arbovirus and a most devastating suid pathogen that, originating from a sylvatic cycle in Africa, has spread to eastern Europe and recently reached western Europe and Asia, leading to a socio-economic crisis of global proportion. However, since neither closely related viruses nor integrated viral elements have yet been identified, ASFV evolution remains a mystery.

**Results:** Here, we show that soft ticks of the *Ornithodoros moubata* group, the natural arthropod vector of ASFV, harbour African swine fever virus-like integrated (ASFLI)-elements corresponding to up to 10% (over 20 kb) of the ASFV genome. Through orthologous dating and molecular clock analyses, we provide data suggesting that integration occurred over 1.47 million years ago. Furthermore, our data indicate that these elements, showing high sequence identities to modern ASFV, are maintained in the tick genome to protect the tick from infection with specific ASFV-strains through RNA interference.

**Conclusion:** We suggest that this mechanism of protection, shaped through many years of co-evolution, is part of an evolutionary virus-vector “arms race”, a finding that has not only high impact on our understanding of the co-evolution of viruses with their hosts but also provides a glimpse into the evolution of ASFV.

## Background

The pandemic spread of viral pathogens affecting humankind has been recognised for many years as one of the most dangerous scenarios leading to crises of global proportion. Furthermore, the “One Health” concept of today is arguing that pathogens affecting animals or plants may also have major impact on the human population (1, 2). One recent example of such interrelations with an extreme global socio-economic impact is the unprecedented pandemic spread of African swine fever virus (ASFV), one of the most devastating viral diseases of animals (3).

ASFV, a double-stranded DNA (dsDNA) virus, can infect all kind of suids and leads to a multi-systemic disease, African swine fever (ASF), in non-natural suid hosts with clinical signs of a viral haemorrhagic fever characterised by morbidity and case fatality rates of up to 100 % (4, 5). Despite almost 100 years of intensive research and the occurrence on four continents (6), neither a vaccine nor any treatment are available. Therefore, the introduction of ASFV into the wild boar populations of eastern Europe in 2007 has led to an unprecedented epidemiological situation that culminated in the introduction of ASFV into western Europe and Asia (7–9). Especially in China, the largest producer of pork in the world, extreme socio-economic consequences can be observed, impacting not only Asia, but affecting the global agriculture and food industry and shifting entire markets (3, 10).

Nevertheless, very little is known about ASFV evolution. ASFV appears to be the only member of its genus (*Asfivirus*) and family (*Asfarviridae*) and is the only known DNA arbovirus (4). First described from, and endemic in, sub-Saharan Africa (11), ASFV is transmitted in a sylvatic cycle between soft ticks of the *Ornithodoros moubata* complex and indigenous wild pigs such as African warthogs (*Phacochoerus africanus*) (12). Both groups of natural hosts are well adapted to an ASFV infection and, although persistently infected (12), show no obvious pathological signs, which has led to the hypothesis that - similar to other arboviruses - ASFV and its hosts have undergone a long time of co-evolution. Since viruses leave no fossil records, analyses into the evolution of ASFV have been restricted to highly similar genotypes of which only very few whole genome sequences are available (4, 13). However, with the recent advances in sequencing technologies, novel ways of analysing virus evolution have been discovered, with one of those being the analysis of endogenous viral elements (EVE) - ancient viral sequences integrated into the host genome (14).

In our study, we identified African swine fever virus-like integrated (ASFLI)-elements in the uncharacterised genome of *O. moubata* complex soft ticks. We show ASFLI-elements in tick cell lines (15), *O. moubata* and *O. porcinus* soft ticks, and in museum-stored *Ornithodoros* ticks collected about 100 years ago in the field in East Africa, allowing for phylogenetic reconstruction and orthologous dating. To evaluate whether the ASFLI-elements retained a function, we conducted tick infection experiments, transcriptome analyses and small RNA sequencing and expressed a highly conserved ASFV-like gene from the tick genome for further characterisation.

## Results

### Evidence of ASFLI-elements in the *O. moubata* tick cell genome

In order to test for the presence of either novel DNA-viruses or integrated viral elements, we sequenced DNA from the *O. moubata* cell line OME/CTVM21 (15), and combined reads from different sequencing platforms. In detail, 14.9 million paired-end reads with 300 bp read length (Illumina MiSeq) and 277 million single-end reads with 50 bp read length (Illumina HiSeq) were combined with about 500,000 3^rd^ generation sequencing single molecule, ultra-long reads (MinION) (Supplementary Table 1).

Using the SPAdes software (16) for the assembly and BLASTn for the detection of viral sequences, we identified 34 contigs with lengths between 336 and 85582 bp containing ASFLI-elements (Figure 1 and Supplementary Tables 2, 3 and 4). However, we detected truncated ASFV-like ORFs as well as sequence duplications and reorganisations when comparing ASFLI-elements with the homologous ASFV genes and the ASFV genome organisation (Figures 1 and 2 and Supplementary Tables 3 and 4). For validation of the assembly, we confirmed two large ASFLI-elements (6.8 kb and 4.2 kb) by PCR and subsequent sequencing (Illumina MiSeq) (Figure 1). The BLASTn search as well as the analysis of the >500 bp ORFs by BLASTp resulted in the identification of contigs showing both integrated viral elements and sequences with similarity to the genome of the black-legged tick *Ixodes scapularis*, the only tick species with a fully sequenced and partially-annotated genome (17). Moreover, contigs were identified that cover multiple ORFs with similarity to mobile genetic elements (Figure 1 and Supplementary Tables 3 and 4). Altogether, we were able to identify the first ASFLI-elements homologous to 46 ASFV-genes, covering 14 genes completely and 32 genes partially, with identities to recent ASFV sequences of up to 86.6 % (Figure 2 and Supplementary Tables 3 and 4).

**Figure 1.**
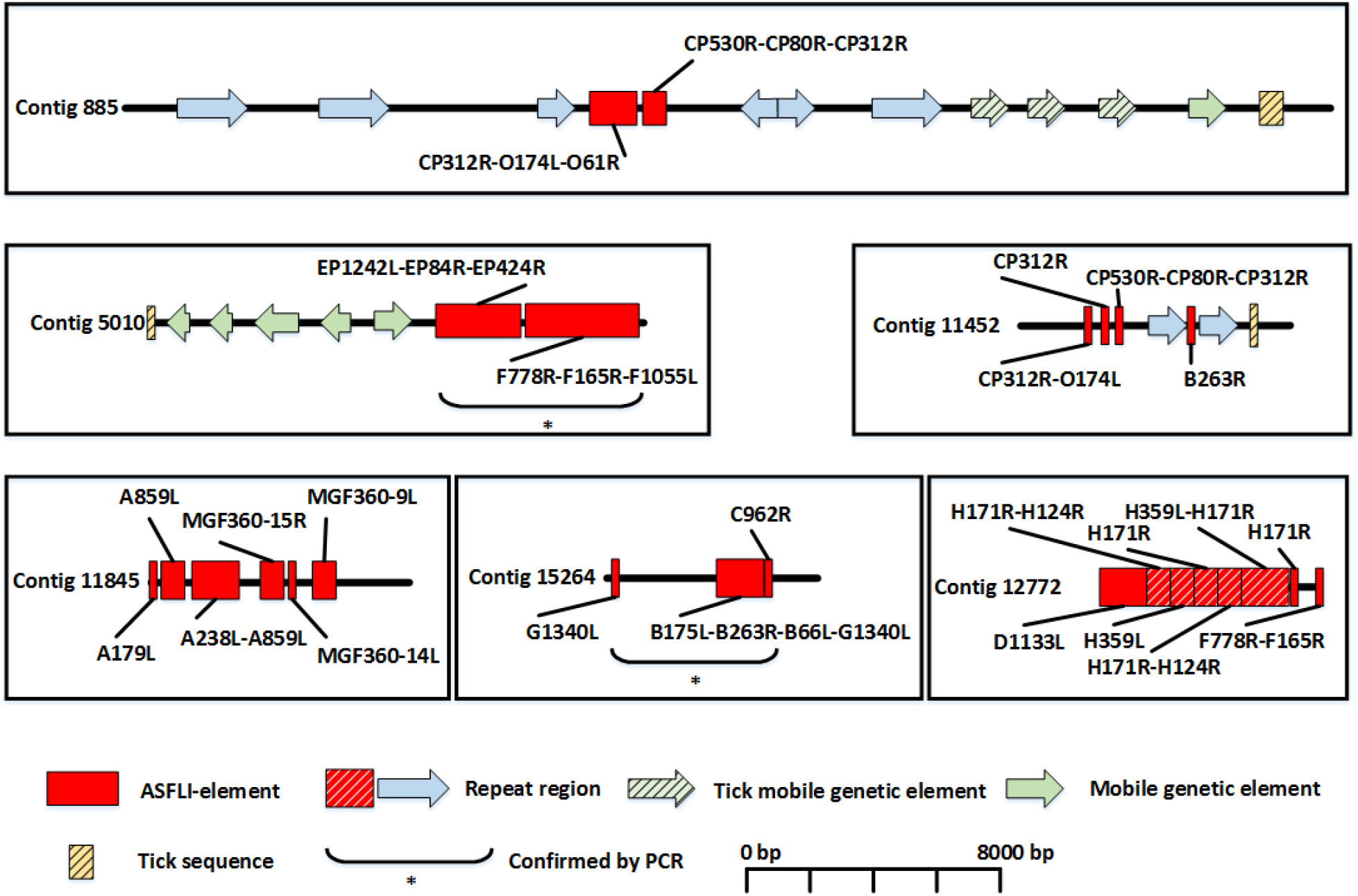
Integration sites of ASFLI-elements in the genome of the *Ornithodoros moubata* cell line OME/CTVM21 genome. SPAdes-assembled contigs showing ASFLI-elements (red) adjacent to tick sequences (yellow-striped), genes from mobile genetic elements (green and green striped arrows) and repeat regions (blue arrow, red-striped) identified by BLASTn and BLASTp. PCR and subsequent sequencing validated the assembly for two major ASFLI-elements (*).

**Figure 2.**
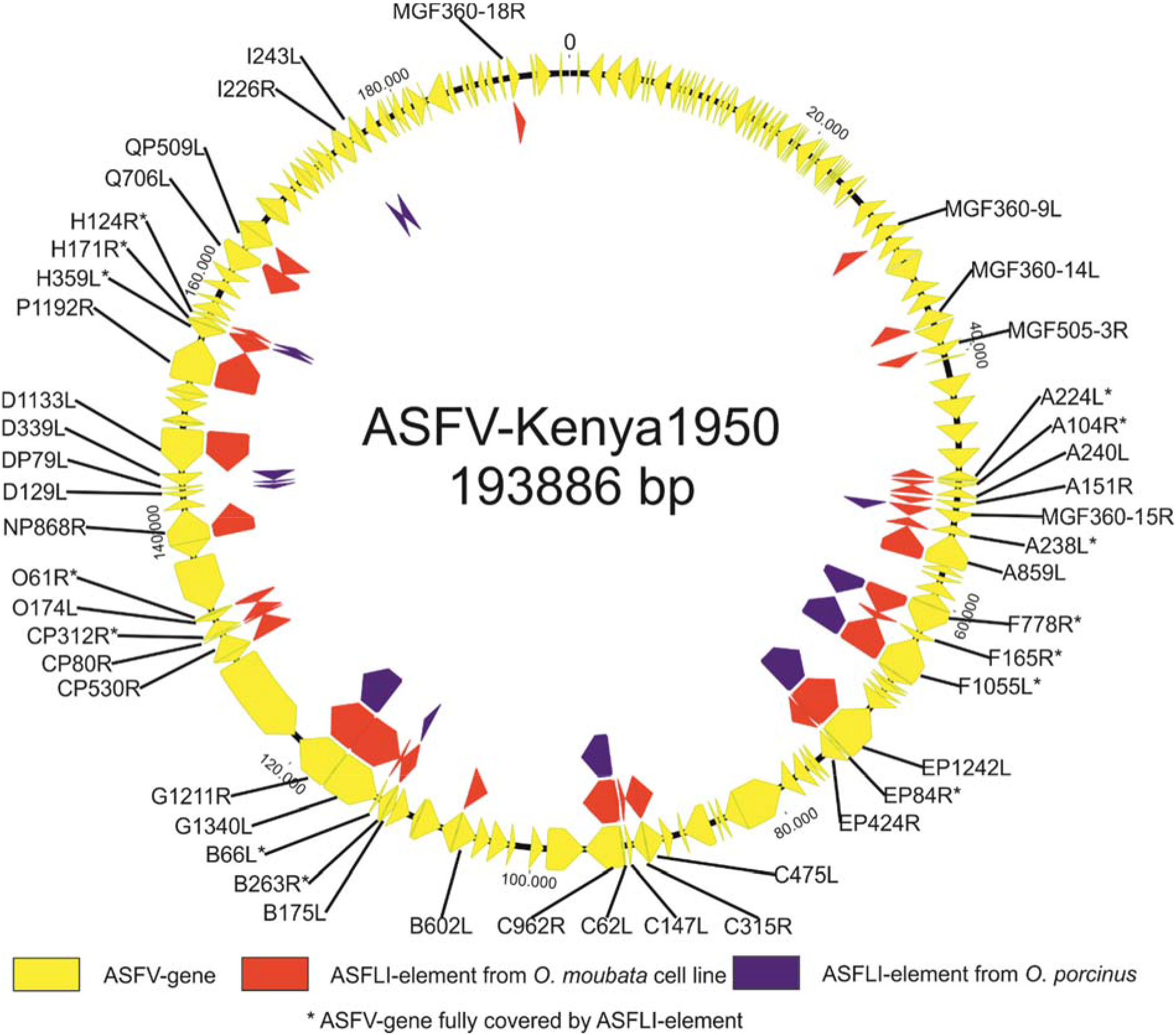
Alignment of ASFLI-elements and homologous ASFV-genes detected in Ornithodoros *moubata* tick cell lines and *Ornithodoros porcinus* ticks. ASFLI elements from *O. moubata* (red) and *O. porcinus* (blue) were identified by BLASTn through their homologies to 32 partial and 14 complete (*) ASFV genes (yellow) distributed over the entire ASFV-Kenya 1950 (Acc. No. AY161360) core genome. For better visualisation, the viral genome is displayed as a circle.

### Phylogenetic analysis shows ASFLI-elements are basal to all known ASFV sequences

In order to place the ASFLI-elements in the context of other nucleocytoplasmic large DNA viruses (NCLDV), the ASFLI-elements corresponding to ASFV proteins pF1055L (Helicase) and pEP1242L (RNA polymerase subunit 2), were aligned with homologous NCLDV amino acid sequences including sequences of ASFV (Supplementary Appendix 1). The amino acid sequence trees of the NCLDV are diverse, but both ASFLI-elements, pF1055L (Figure 3) and pEP1242L (Supplementary Appendix 1), group with the ASFV amino acid sequences, thus forming well-supported monophyletic clades (bootstrap value = 1). The ASFLI-elements are, however, placed as a close outgroup to the other ASFV sequences (Figure 3 and Supplementary Appendix 1).

**Figure 3.**
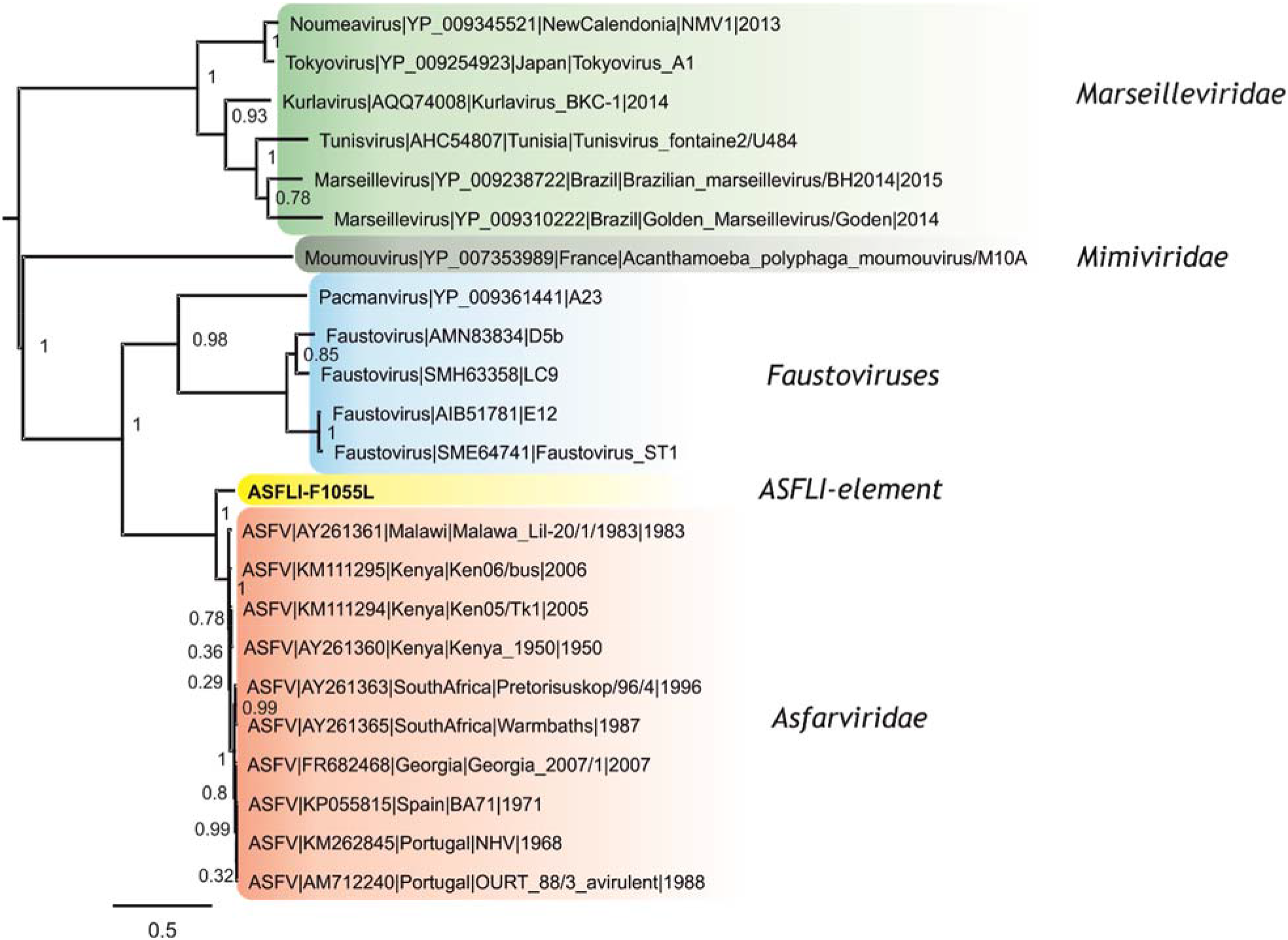
Phylogeny of ASFLI-pF1055L and ASFV homologues. Maximum likelihood tree (JTT+G_4_) showing ASFLI-element ASFV-sequences and NCLDV background protein sequences for F1055L (Helicase) and analogues. Statistical support of 100 bootstraps is indicated at the nodes.

### ASFLI-elements are present in recently-sampled *O. moubata*, *O. porcinus* and approx. 100-year-old *O. moubata* and *O. porcinus* field-collected ticks from Africa

To evaluate if the integration of the ASFLI-elements is specific for the *O. moubata* cell line OME/CTVM21, we tested all six available *Ornithodoros* cell lines as well as laboratory-reared *O. moubata* ticks from various origins using NGS and quantitative PCR (qPCR) for ASFLI-elements. Furthermore, we analysed *O. porcinus ticks* collected from the field in Kenya in 2017 and from a French laboratory-reared colony, *Ornithodoros erraticus* field ticks from Portugal and *Ornithodoros savignyi* from Nigeria. While all tick cell lines and the *O. moubata* and *O. porcinus* ticks tested positive for ASFLI-elements, *O. savignyi* and *O. erraticus* (Supplementary Table 7) tested negative. By sequencing DNA from the *O. porcinus* ticks (Supplementary Table 1) and assembly, we were able to identify 18 contigs containing ASFLI-elements with lengths over 150 bp that are related to 14 ASFV genes (Figure 2 and Supplementary Tables 1 and 5). For five of these genes, corresponding ASFLI-elements were only detected in *O. porcinus* (Figure 2 and Supplementary Table 1 and 5). One orthologous element, including the ASFV-like EP1242L gene, was discovered in the same insertion site in *O. moubata* and *O. porcinus* (98.5 % nucleotide sequence identity flanking a piggyBac transposable element with 90.3 % nucleotide sequence identity) (Supplementary Figure 6).

Through the analysis of published transcriptome data from *Ornithodoros* ticks, we identified ASFLI-specific RNA in *O. moubata* ticks from laboratory colonies from Spain (18) and Japan (19) (Figure 4 and Supplementary Tables 1 and 2), while in pertinent data from *O. erraticus (20)* and *Ornithodoros turicata* (PRJNA447876) no ASFLI-specific RNA was detected (Figure 4). Furthermore, we examined nine *Ornithodoros* specimens collected between 1906 and 1913 in the former German colonies of East and southern Africa, that had been stored in ethanol at the Berlin Museum for Natural History (MfN, by sequencing (Figure 4 and Supplementary Tables 1 and 6). In all libraries relating to *O. moubata* and *O. porcinus* ticks, we detected ASFLI-specific reads or even contigs (Figure 4, Supplementary Table 6).

**Figure 4.**
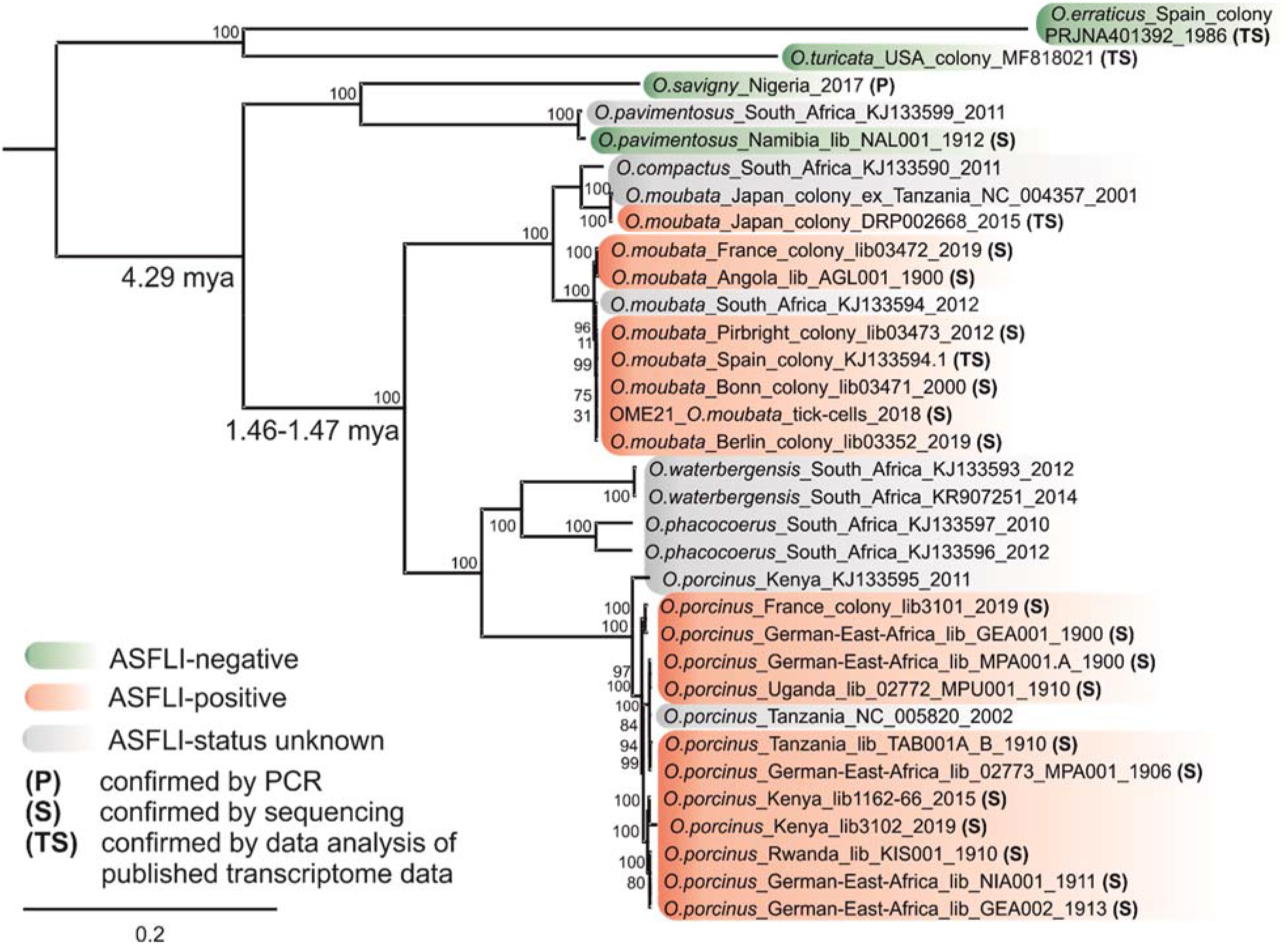
*Ornithodoros* phylogeny and orthologous dating of ASFLI-element integration. A maximum-likelihood (ML) tree was constructed using IQ-TREE v1.6.5 with standard model selection, resulting in the best-fit model TIM2+F+R3 (AC=AT, CG=GT and unequal base frequencies + empirical base frequencies + FreeRate model with 3 categories) based on MAFFT v7.388 aligned full-length mitochondrial sequences from *Ornithodoros* ticks. Statistical support of 10,000 ultrafast bootstraps is indicated at the nodes. Taxon names include, where available, tick species designation, country of origin, library number, INSDC accession number and sampling date. Clock data on tick species divergence was included from the literature (21).

### Phylogenetic reconstruction using full-length mitochondrial genomes of soft ticks reveals ASFLI-integration might have occurred over 1.46-1.47 million years ago (mya)

To assess the phylogenetic relationship between the different tick species carrying ASFLI-elements and perform orthologous dating, we generated full-length mitochondrial genomes from the tick sequence data generated in this study (Supplementary Table 1). These genomes were aligned together with additional tick mitochondrial genome sequences from the literature (21) using MAFFT v7.388 in Geneious, and a phylogenetic tree (Figure 4) was constructed using IQ-TREE v1.6.5. Molecular clock data from the literature on tick species divergence, that was calibrated using fossil data (21), suggests that the *O. moubata* group including *O. moubata*, *O. porcinus*, *O. compactus*, *O. waterbergensis* and *O. phacocoerus* originated 4.29 mya and that *O. moubata* and *O. porcinus* probably diverged 1.46-1.47 mya (21). Since we detected an orthologous ASFLI-element in both *O. moubata* and *O. porcinus* (Supplementary Figure 6) suggesting at least one joined integration event, we can estimate a minimum time to the integration of this ASFLI-element to be 1.46-1.47 mya (Figure 4). Because we did not discover ASFLI-elements in ticks outside the *O. moubata* group, e.g. *O. erraticus, O. turicata, O. savigny and O. pavimentosus* (Figure 4), we can further estimate a maximum time of the integration of 4.29 mya.

### Molecular clock analyses using ASFLI-elements from different *Ornithodoros* species provide an estimate for a time to the most recent common ancestor consistent with orthologous dating

In addition to the mitochondrial genomes, we analysed the orthologous ASFLI-element containing ASFV-EP1242L from five soft tick specimens (three *O. moubata* and two *O. porcinus*). These sequences contained deletions and one insertion in comparison to the homologous ASFV-EP1242L. Maximum likelihood phylogenies from the corresponding amino acid sequences revealed that all samples clustered as an outgroup close to the *Asfarviridae* (Supplementary Appendix 1). Due to missing data points over time that would be useful for calibration, inferring time-scaled phylogenies on the EP1242L containing ASFLI-element sequences from the soft tick samples was more challenging than for the mitochondrial genes. However, by using a strict clock with molecular clock rate priors of 5e-7 to 1e-8 substitutions per site per year (commensurate with the expected substitution rate for host DNA), a time to most recent common ancestor (TMRCA) of 0.08 to 4.7 million years was inferred (Supplementary Appendix 1). Although the confidence intervals on these TMRCA are very wide, the TMRCA of the EP1242L ASFLI-element is consistent with the overall integration estimate of 1.46 – 4.29 million years obtained from orthologous dating using the host mitochondrial genomes (Supplementary Appendix 1).

### ASFLI-elements may influence ASFV-infection in ticks and tick cell lines

In order to investigate a possible influence of the identified ASFLI-elements on ASFV infection of soft ticks, we infected *O. porcinus* ticks from Kenya and *O. moubata* ticks from the Berlin colony by membrane-feeding with 1 × 10^4^ and 1 × 10^6^ haemadsorbing units (HAU)/ml of the ASFV ken.rie1 P72 genotype × strain that was originally isolated from a Kenyan *O. porcinus* tick. To analyse the influence of different ASFV-genotypes on the infection rate of the ticks, *O. porcinus* and *O. moubata* ticks were furthermore infected with 1 × 10^5^ HAU/m of an ASFV-Ken06.bus genotype IX strain or with 1 × 10^4^ HAU/ml of an ASFV-Sardinia genotype I isolate. All tick specimens were tested for the accumulation of the late-expressed *p72* specific viral transcripts by quantitative reverse-transcriptase PCR (RT-qPCR), and the virus titre of the blood-virus solution used for oral infection was determined by titration on porcine macrophages.

In the initial experiment (Figures 5 A and C), all *O. porcinus* ticks (25/25) showed a clear accumulation of viral RNA while from the *O. moubata* ticks, only 5 % (2/40) accumulated viral transcripts after infection with the higher virus dose (Figures 5 B and D and Supplementary Table 9). While among the ticks infected with ASFV Ken06.bus, a minimal *p72* transcript accumulation was detected only in *O. moubata* (3/25) (Figures 5 E and F), four *O. moubata* (4/25) and three *O. porcinus* ticks (3/10) infected with ASFV Sardinia showed a minimal increase of *p72* transcripts (Figures 5 G and H). Furthermore, infection of the tick cell lines OME/CTVM21, OME/CTVM22, OME/CTVM24 and OME/CTVM27 with ASFV ken.rie1 did not result in any detectable viral transcription (data not shown).

**Figure 5.**
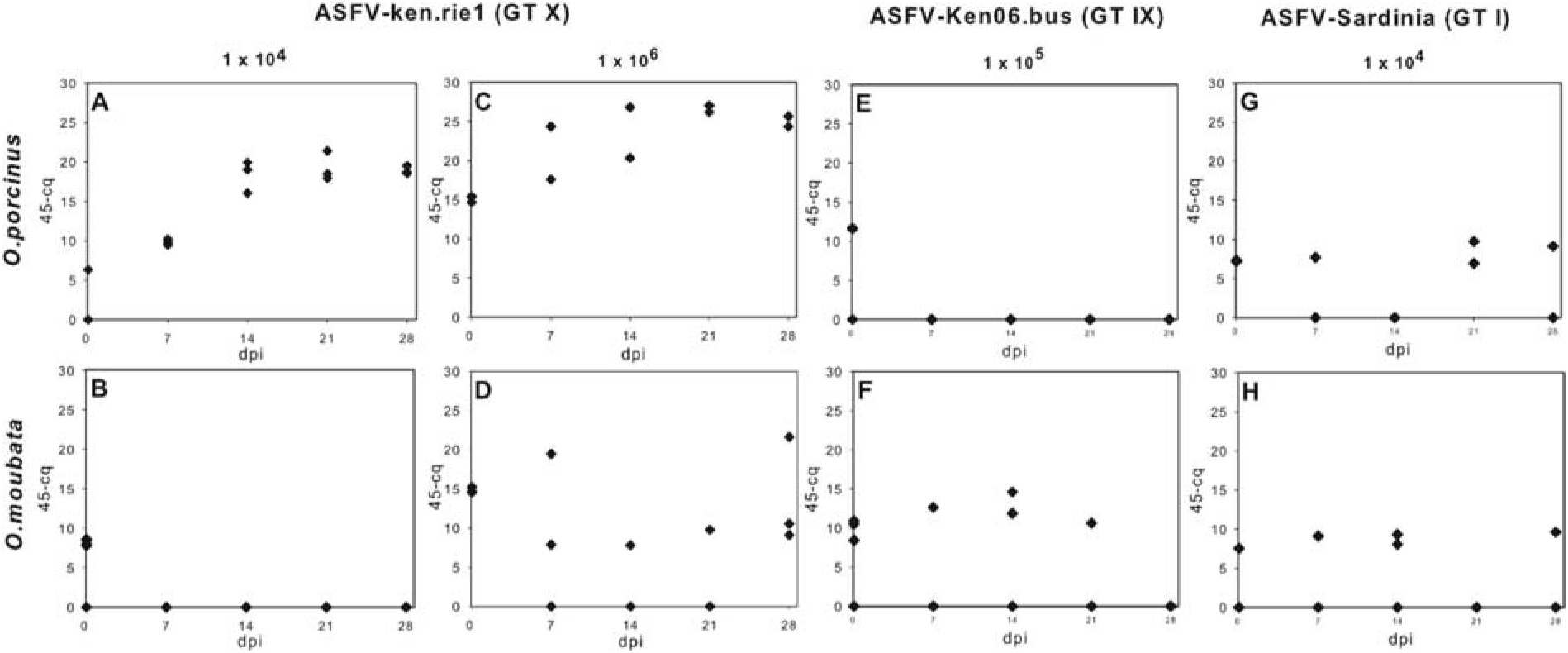
RT-qPCR results of *Ornithodoros spp.* ticks experimentally infected with different ASFV genotypes. Shown are ASFV-P72 transcript-specific Cq-values of third nymphal stage ticks fed with defibrinated pig blood containing either 1 × 10^4^ HAU/ml or 1 × 10^6^ HAU/ml ASFV-ken.rie1 (GT X) **(A-D)**, 1 × 10^5^ HAU/ml ASFV-Ken06.bus (GT IX) **(E-F)** or 1 × 10^4^ HAU/ml ASFV-Sardinia (GT I) **(G-H).** Due to the limited number of field ticks available and feeding under artificial conditions, fifteen *O. porcinus* ticks were collected in each of three experiments **(A, E)** and ten in two experiments **(C, G)** while for the laboratory-reared *O. moubata,* twenty-five individuals were collected in each of three experiments **(B, F, H)** and fifteen in one experiment **(D)**.

### RNA sequencing demonstrates ASFLI-specific mRNA - small-interfering and piwi-interacting RNAs in tick cells

Interference with small RNAs (RNAi), e.g. small interfering RNA (siRNA) or piwi-interacting RNA (piRNA), could be responsible for the observed differences in ASFV infection rates of the *Ornithodoros* ticks (as shown in insects (22)); we therefore investigated hypothetical ASFLI-element-specific small RNAs in the ticks and performed small RNA sequencing (Supplementary Figures 1-4 and Tables 1 and 10). The distribution of the read lengths showed a distinct bimodal distribution in all libraries with one peak at 22nt, representing siRNAs, and one at 28-29nt, representing piRNAs (as shown by the typical U-bias at the 5’-end) (Supplementary Table 10, and Supplementary Figure 1) (23). This is in accordance with small RNA sequencing data from hard ticks (24, 25). By mapping the resulting data (siRNA and piRNA fractions) against the ASFLI-element-containing contigs, we identified siRNAs and piRNAs homologous to different ASFLI-elements (Supplementary Figures 2 and 3). While for some elements none or only single homologous siRNA/piRNA molecules were identified, elements related to the ASFV-genes EP1242L and F778R showed multiple corresponding siRNAs/piRNAs.

Following the mapping of all siRNAs and piRNAs against available ASFV whole-genome sequences from different viral genotypes, we observed matching RNAs but identified differences in the number of mapped reads from the different tick species. While the highest number of siRNA and piRNA from *O. moubata* mapped to ASFV genotypes IX and X, piRNA from *O. porcinus* mapped best to ASFV genotypes II, III, IV and V (Figure 6, Supplementary Table 10, Supplementary Figures 2, 3 and 4). The siRNA fraction of *O. porcinus*, however, mapped best to genotype IX and X isolates. In addition to the small RNA, we detected ASFLI-specific mRNA in the OME/CTVM21 cells as well as in *O. moubata* transcriptome data from the literature (Supplementary Tables 1, 2 and 8).

**Figure 6.**
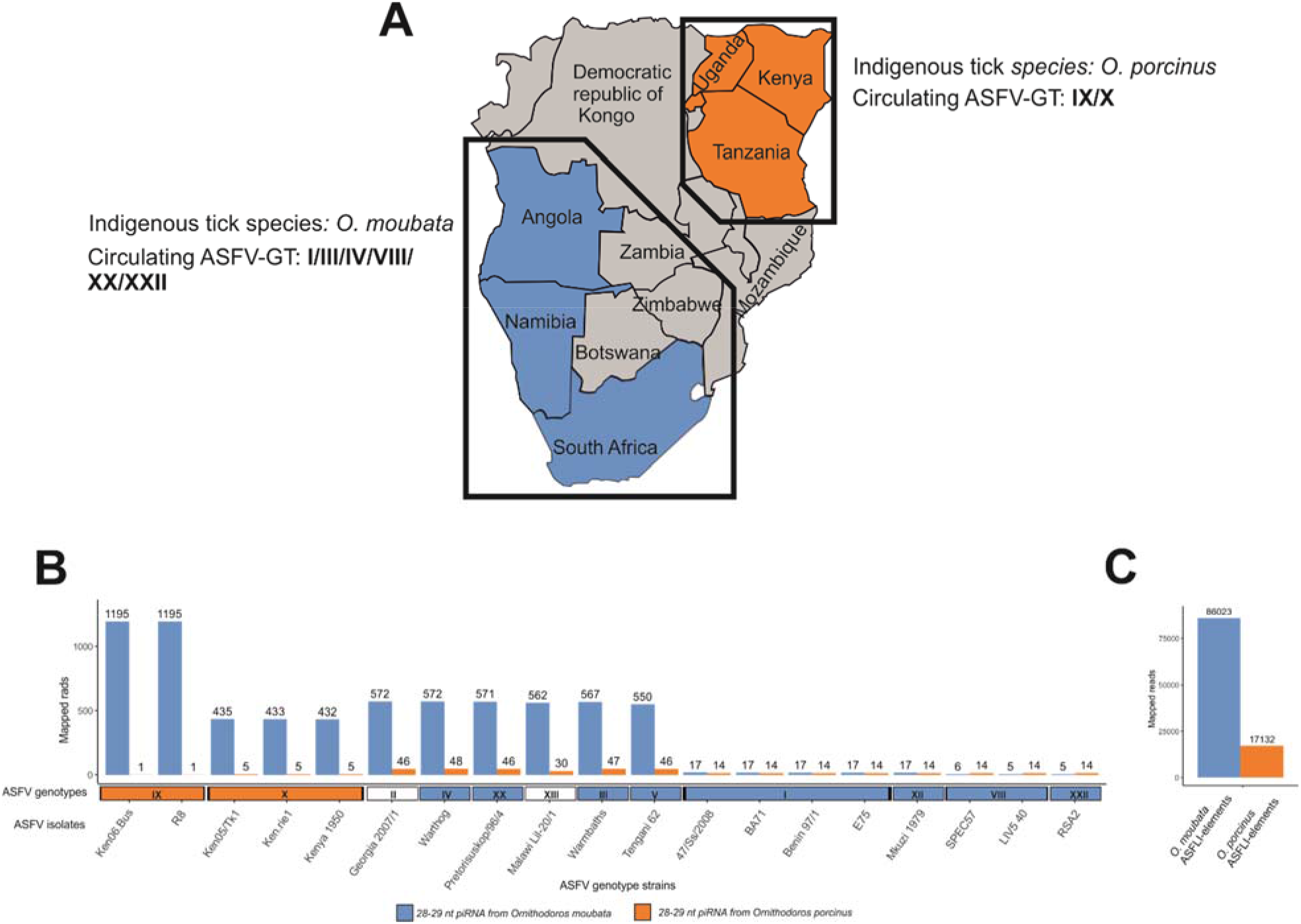
Regional distribution of *Ornithodoros moubata*, *Ornithodoros porcinus* and ASFV-genotype strains, and differences in *O. moubata* and *O. porcinus* piRNA identity with specific ASFV-genotypes and ASFLI-elements. piRNA (28-29nt) reads from *O. moubata* (originating from southern Africa) and *O. porcinus* ticks (originating from East Africa) **(A)** were mapped against available ASFV whole-genome sequences from different genotypes with Bowtie2 **(B)**. While the lowest number of mapped *O. moubata* piRNA reads was observed for ASFV-genotypes shown to infect *O. moubata* (ASFV-GT I, XII, VIII, XXII), the lowest number of *O. porcinus* piRNA reads mapped to ASFV-genotype X able to infect *O. porcinus* **(B)**. With up to 100 % sequence identity, 86.023 (O. moubata) and 17.132 (O. porcinus) reads mapped to the ASFLI-elements **(C)**.

### The reconstructed ASFV-like A104R protein is highly similar to its ASFV homologue but is not expressed in tick cell lines

Among the complete ASFLI-element genes discovered by NGS, ASFLI-A104R is one of the most conserved with nucleotide sequence identities of 83.0-85.0 %, and amino acid sequence identities of 94.0-95.0 %. However, the insertion of an ‘A’ nucleotide in a homopolymer region of eight ‘A’ nucleotides, starting at position 9 of the gene, leads to a frameshift and stop-codon integration (Supplementary Figure 7). Since no small RNA molecules matching ASFLI-A104R were identified, we investigated whether the ASFLI-A104R protein is expressed in the cultured tick cells. We repaired the alA104R-ORF *in silico* by removing the additional ‘A’, expressed the synthetic gene as both a native protein and a flag-tagged variant in *E. coli* and raised rabbit sera against the purified alA104R proteins. Using Western blot analysis, we observed specific signals for the native protein as well as for the flag-tagged alA104R protein, and also for the ASFV-Kenya 1033 protein. However, no ASFV-like A104R specific signals were obtained with protein extracts from the different tick cell lines (Supplementary Figure 7).

## Discussion

We here report the existence of endogenous ASFLI-elements in the genome of *O. moubata* complex soft ticks. The detection of these ancient integrated sequences will allow us, for the first time, to gain deeper insights into the evolution of ASFV, one of the most dangerous animal pathogens known to date, and into the role of endogenous viral elements in the co-evolution of ticks and viruses.

Generally, very little is known about tick genome organisation, or how and how often viral elements become integrated into a tick genome (26) and for what reasons. We identified ORFs related to mobile genetic elements as well as repeat regions and possible target site duplications formed by the integration of mobile genetic elements adjacent to the ASFLI-elements. Therefore, it might be hypothesised that the integration of viral elements into large hyper-variable regions of the tick genome (17, 27–29) is assisted by the interaction with mobile genetic elements as discussed for endogenous Borna-like N (EBLN)-elements (30) and various arthropod viruses (27, 29, 31, 32).

Further analyses showed that *O. moubata* and *O. porcinus* ticks, in addition to the *O. moubata* tick cell lines, harbour ASFLI-elements. This observation clearly demonstrates the fixation of these elements in the *O. moubata* and *O. porcinus* genomes. While it remains unclear how the fixation occurred, the initial event leading to germ line integration might be explained by the fact that ASFV infects the tick’s reproductive tissues in order to become transovarially and venereally transmitted. Furthermore, although ASFV replication takes place in the cytoplasm, viral mRNAs were demonstrated to enter the nucleus of infected Vero cells and primary porcine macrophages (33, 34), where they could have been integrated.

Although we detected matching ASFLI-elements in different *Ornithodoros* tick species, the occurrence of multiple integration events over time cannot be ruled out. However, the identification of an orthologous ASFLI-element including the ASFV-like EP1242L gene in both *O. moubata* and *O. porcinus* and the analysis of indel-patterns leads to the conclusion that one integration event might have taken place over 1.46-1.47 million years ago. Since we were not able to detect ASFLI-elements in *Ornithodoros* species other than *O. moubata* and *O. porcinus*, we can further estimate a maximum time to the integration of 4.29 million years ago, when these species diverged. Although the molecular clock analysis using the orthologous ASFLI-elements resulted in a TMRCA consistent in range with the orthologous dating, missing data through time hindered calibration and thus, model selection was difficult and the confidence intervals are very wide. Therefore, and since the pattern and rate of substitutions of the ASFLI-elements could have been furthermore influenced by positive selection due to the protective effect against ASFV, the results should be interpreted with caution and more data on ASFLI-elements in different *Ornithodoros* tick species is needed to define the possible time-point of integration more precisely.

In total, we identified ASFLI-elements in the genomes of *O. moubata* and *O. porcinus* ticks homologous to 46 ASFV-genes; most of them represent genes with properties linked to viral DNA replication or RNA transcription. These observations may be explained by the fact that these genes in particular are highly conserved between viral isolates (90-100 %); the genes coding for structural proteins are more variable, therefore we cannot exclude that the latter, if also integrated, are simply not detectable by identity-based search methods like BLAST. However, the integration of major viral structure protein genes might have been harmful for the tick cell, so that they were just not maintained in the genome. Furthermore, it might be possible that that the integration of the conserved viral core genes, essential for the early phases of viral infection, is in particular linked to a beneficial function, such as protection by RNAi.

During infection experiments using *O. moubata*, *O. porcinus* and different ASFV genotypes, we observed differences in the ability of ASFV to infect the various *Ornithodoros* species. While a genotype X isolate from Kenya was able to replicate in *O. porcinus* ticks from Kenya, no infection was observed in *O. moubata* (phylogenetically closest to southern African ticks). However, when using a higher viral titre, successful infection could be observed in a small number of *O. moubata* ticks indicating that, in principle, infection is possible and that low-level infection might be blocked by a tick defence mechanism, such as RNAi.

Through the identification of ASFLI-specific siRNA and piRNA in the ticks and tick cells, we provide data suggesting that the ASFLI-elements might in fact represent an RNAi-based defence against ASFV infection in *Ornithodoros* ticks. Both siRNA and piRNA are known as major regulation factors of transcription, transposon suppression and defence against viral pathogens (35–38). Although, for ticks the known arsenal of antiviral mechanisms includes protection by RNAi (39), the role of EVEs in RNAi and protection against viral pathogens has never been reported.

Whether the integration of non-retroviral sequences into a host genome is a directed process and how often it leads to improved fitness, is controversially discussed (29, 40, 41). However, protection against homologous viruses was demonstrated after integration of retroviral-elements in mice, sheep (41) and bats (27), and is also discussed for EBLN-elements (27, 30). Furthermore, transcriptional activity of EVEs and protection against viral infection was demonstrated to occur in crustaceans (29, 31, 32) and mosquitoes (42, 43).

In addition to the hypothesised protective function, our results might suggest that the ASFLI-element-based protection could be selective for specific ASFV genotypes and therefore different in *O. porcinus* and *O. moubata*. This observation is supported by data from the literature in which clear differences were observed in tick infection rates using different ASFV genotype isolates and tick species (44–46). Our successful infection experiments with a genotype X virus using the *O. porcinus* ticks from Kenya and experiments with the Pirbright and French *O. moubata* colonies and genotype I viruses (46–48) further strengthen this hypothesis.

Therefore, it could be speculated that during millions of years of co-evolution, where ASFV hosts were regionally isolated by geographical factors (e.g. mountains, rivers), ASFV and its soft tick hosts have adapted to co-exist as displayed by the regional distribution of ASFV-genotypes and the lack of any pathological findings in infected soft ticks, despite generalised virus infection and high viral titres. However, additional immunological or genetic factors determining which ASFV strain can infect a specific *Ornithodoros* tick species need to be considered, and more infection experiments using different *Ornithodoros* tick species and ASFV genotypes and data on small RNA are needed to substantiate this hypothesis.

The possible role of EVEs in the antiviral response of ticks might also explain why some EVEs have been protected from accumulating mutations. However, it remains unclear why for the highly identical ASFLI-A104R protein, whose viral counterpart is crucial for virus replication (49), that is one of the most abundant ASFV proteins in infected wild boar lung cells and infected Vero cells (50) and is highly conserved between ASFV isolates (49), no siRNA or piRNA could be detected. While our data might lead to the conclusion that the ASFLI-A104R protein is not translated, the amount of protein might just have been too low for detection or cellular machinery might degrade the protein. Due to the high identity to modern ASFV-A104R sequences, a more recent integration event has also to be considered. However, further analyses including mass spectrometry analyses, the cloning of the ASFLI-A104R element into a recent ASFV isolate and the biochemical and functional characterisation of the protein should be performed to elucidate a possible function and the nature of ASFLI-A104R.

## Conclusion

In conclusion, we present evidence for endogenous viral elements of the only known DNA arbovirus, ASFV, which might have been integrated into soft tick genomes over 1.46-1.47 million years ago. They serve as templates for siRNA and piRNA thereby possibly protecting the tick against viral infection. We believe that this discovery is an exceptional starting point to study the evolution of ASFV, one of the most dangerous threats to sustainable pig production in Africa, Europe and Asia, as well as the mechanisms by which organisms adapt to the ancient struggle with viral pathogens.

## Materials and Methods

### Virus strains

For infection studies, we used an ASFV isolate designated “ASFV ken.rie1”. The virus was isolated from *O. porcinus* ticks from Kenya on swine peripheral blood mononuclear cells (PBMCs), passaged twice and titrated on PBMCs as described elsewhere (51). By sequencing the *p72*, *B602L* and *p54* genes, we compared “ASFV ken.rie1” to known ASFV isolates from GenBank and grouped it into *p72* genotype X, which is known to circulate in East Africa between ticks and indigenous pigs (52). “ASFV-Sardinia” and “ASFV ken06.bus” (originally received from the European reference laboratory for ASFV (CISA-INIA)) were stored at the German national reference laboratory for ASFV as reference material. Both had initially been passaged and titrated on PBMCs.

### Tick rearing, tick infection and tick cell cultures

*O. moubata* ticks, sourced from laboratory colonies maintained at HU-Berlin, Germany, and CIRAD-France as well as *O. porcinus* ticks collected from the field in Kenya and *O. erraticus* ticks collected from the field in Portugal were held at 25 °C and 85 % relative humidity in the dark and fed on defibrinated pig blood as described elsewhere (53). Laboratory-reared *O. moubata* from The Spanish National Research Council (CSIC) Salamanca in Spain and The Pirbright Institute in the United Kingdom were received and stored in 70% ethanol. DNA from Nigerian *O. savigny* was provided by the University of East London. Tick infection was carried out using the routine feeding system with ASFV-spiked blood. The *O. moubata* cell lines OME/CTVM21, OME/CTVM22, OME/CTVM24, OME/CTVM25, OME/CTVM26 and OME/CTVM27 were maintained in sealed flat-sided culture tubes (Nunc, Thermo Scientific, Waltham, Massachusetts) at 28 °C as described elsewhere (15). Tick cells were infected with ASFV by adding 0.22 × 10^5^ HAU directly to 2.2 ml cell culture without washing. On days 1-7, 14, 21 and 28 post infection, 125 μl cell culture were removed and added to 125 μl PBS and 750 μl TRIzol LS (Thermo Scientific) for RNA extraction. An equal volume of fresh culture medium was added to the culture tubes. All samples were stored at 4 °C until RNA extraction.

### Nucleic acid extraction

Ticks were homogenised in a 2 ml reaction tube with two 5 mm steel beads and 500 μl PBS using a Tissue Lyser II (Qiagen, Hilden, Germany) for 3 min at 30 Hz. The homogenate was centrifuged at 10,000 × *g* for 5 min, and DNA was extracted from the supernatant using the High Pure Template Prep Kit (Roche) according to the manufacturer’s instructions. Tick cell line DNA was extracted using the same kit according to the manufacturer’s instructions.

High molecular weight DNA was isolated from tick cells using a salting-out protocol (54), slightly modified by adding RNase A (10 mg/ml) for incubation at 37 °C for 60 min in the first step and proteinase K (10 mg/ml) for incubation at 56 °C for another 60 min in the second step.

For RNA extraction, ticks were homogenised in a 2 ml reaction tube with two 5 mm steel beads and 1 ml TRIzol Reagent (Thermo Scientific) using a Tissue Lyser II (Qiagen) for 3 min at 30 Hz. For efficient cell lysis, the homogenate was incubated for 10 min at room temperature. After addition of 200 μl trichlormethane (Sigma Aldrich, St. Louis, Missouri), samples were thoroughly mixed, incubated for 10 min at room temperature and centrifuged at 10,000 × *g* for 10 min at 4 °C. The clear aqueous phase was removed, mixed with 600 μl of 100 % ethanol and loaded onto an RNeasy-spin column (Qiagen) for RNA extraction and additional on-column DNA digestion following the manufacturer’s instructions. RNA extraction from tick cell lines was performed using TRIzol LS and RNeasy-spin columns (Qiagen) according to the manufacturer’s protocol. Prior to qRT-PCR analysis, all RNA samples were treated with Turbo-DNase (Life Technologies) for removal of residual DNA.

DNA extraction from museum-stored ticks was done in a DNA cleanroom under ancient-DNA (aDNA) conditions as described elsewhere (55, 56).

### Oligonucleotide design

All primers and probes were designed using the Geneious (v.10.0.9) software suite and are listed in Supplementary Table 12 with additional primer sequences obtained from the literature.

### PCR

For PCR reactions, Phusion Green Hot Start II High Fidelity PCR Master Mix (Thermo Scientific) was used according to the manufacturer’s instructions on a C1000 Thermo Cycler (Bio-Rad, Hercules, California). PCR products were visualised, depending on their length, on 1 % or 1.5 % agarose gels and visualised by ethidium bromide staining under UV light.

### qPCR and RT-qPCR

All qPCR and RT-qPCR-reactions were done using the QuantiTect Multiplex PCR NoROX Kit (Qiagen) or QuantiTect Probe RT-PCR Kit (Qiagen) in a 12.5 μl reaction including 2.5 μl extracted DNA or RNA, on a C1000 Thermo Cycler (Bio-Rad) in combination with an CFX96™ Real-Time PCR Detection System (Bio-Rad) according to the manufacturer’s instructions. Primers and probes were premixed (10 pmol primer/μl + 1.25 pmol probe/μl), and 1 μl of the mixture was used per reaction. For ASFV DNA amplification, an OIE listed qPCR assay was used targeting the viral B646L gene as described elsewhere (57). For RT-qPCR analysis of ASFV transcripts, the same OIE-listed qPCR was used, only modified by using the QuantiTect Probe RT-PCR Kit (Qiagen).

Control qPCR systems targeted *O. moubata* mitochondrial (16S rDNA) modified from (58), *O. moubata* chromosomal (enolase) and *O. erraticus* chromosomal (subolesin) housekeeping genes (designed in this study). For all samples tested by RT-qPCR, controls without reverse transcriptase were included to demonstrate the absence of DNA. For absolute quantification, a standard curve was generated, using samples of known copy number produced from cloned gene fragments, and measured by the Quant-iT dsDNA Assay Kit HS (Thermo Fisher Scientific).

### Sanger sequencing

For Sanger sequencing, PCR products were extracted from the agarose gel with the QIAQuick Gel Extraction Kit (Qiagen). The sequencing-PCR was done using the BigDye Terminator Kit V1.1 (Thermo Scientific). After purification with NucleoSEQ spin columns (Macherey Nagel, Düren, Germany), samples were sequenced on a Genetic Analyzer 3031XL (Life Technologies, Carlsbad, California), followed by chromatogram editing with Geneious (v.10.0.9) software.

### Next-generation sequencing

For IonTorrent PGM sequencing, the libraries lib01543-lib01546, lib01610 and lib01611 were prepared from the six *O. moubata* cell lines as detailed in Supplementary Table 1 and sequenced as described elsewhere (59). The libraries from museum-stored ticks (lib03076-3086) were prepared under aDNA conditions in a DNA cleanroom and sequenced as described elsewhere (55, 56).

The libraries Lib02151 and Lib02152 were prepared from OME/CTVM21 cells for sequencing on an Illumina MiSeq platform in 300 bp paired-end mode as previously described (60) (Table 1). The library Lib02339 was prepared from OME/CTVM21 cells for sequencing on an Illumina HiSeq platform in 50 bp paired-end mode (Supplementary Table 1). Extracted DNA was fragmented to a size of 150 bp (M220 Focused-ultrasonicator™, Covaris, Woburn, Massachusetts), and libraries were prepared using NEXTflex™ Dual-Index adapters (Bioo Scientific, Austin, Texas) in an automated fashion (SPRIworks Fragment Library System II, Beckman Coulter, Brea, California). Size exclusion was performed manually aiming at a fragment length peak of 350 bp (Agencourt® AMPure® XP Beads, Beckman Coulter). The library was then amplified (AccuPrime™ Taq DNA Polymerase, high fidelity, Invitrogen, Carlsbad, California), purified (Agencourt® AMPure® XP Beads) and quality-checked (High Sensitivity DNA Kit, Agilent Technologies). The final library was quantified (KAPA Library Quantification Kit, Roche) and sequenced (Illumina) according to the manufacturer’s instructions.

For MinION sequencing, high molecular weight tick cell DNA was fragmented using G-Tubes (Covaris), followed by library preparation (LibMinION_1) using the 1D^2^ Sequencing Kit (R9.5) (Oxford Nanopore, Oxford, United Kingdom) according to the manufacturer’s protocol. Sequencing was performed on the MinION MIN-101B system (Oxford Nanopore).

For small RNA sequencing, library preparation was performed using the Ion Total RNA-Seq Kit v2 (Thermo Scientific). The libraries lib02965 (mRNA), lib02966, lib02967, lib03460 and lib03506 (small RNA) were prepared for, and sequenced on, an IonTorrent S5 (Thermo Fisher Scientific) as described elsewhere (61).

### Data analysis

The metagenomic classification software pipeline RIEMS (62) was used on Lib01543-01546 and Lib01610-01611 (Supplementary Table 1). For further data analysis and proof of integration of ASFV-like sequences into the tick cell genome, overlapping read pairs in the MiSeq data sets (Lib02151 and Lib02152) were merged to single reads using FLASH v1.2.11 (63). Subsequently, the merged reads were assembled together with data from Lib02339 and LibMinION_1 using SPAdes v3.11.1 (16) in the mode of read error correction prior to assembly. In total, 66,745 contigs were assembled with standard assembly parameters. The resulting contigs were blasted (BLASTn, NCBI, v2.6.0+) against a customised database comprising all sequences with the NCBI taxonomy ID 10497 (African swine fever virus) (as of 16 January 2018). Hits were filtered using a cut off e-value of 1×10^−4^ and a minimum alignment length of 150 bp, resulting in 34 contigs. These were then blasted against the complete NCBI database to reliably identify and annotate ASFV-like sequences and areas of the host genome.

ASFLI-element sequences from mRNA sequencing data and additional *O. moubata* transcriptome data (18) downloaded from Genbank were identified by mapping using the Newbler software 3.0 (Roche) against a database containing all known ASFV-sequences and previously-obtained ASFLI-element sequences.

Data from the *O. porcinus* ticks were assembled using SPAdes v3.11.1 (16) in the mode of ‘read error correction prior to assembly’. The resulting contigs were blasted against a customised database comprising all sequences with the NCBI taxonomy ID 10497 (African swine fever virus) (as of 10 May 2019). Obtained hits were filtered as described above.

Data from the museum ticks were mapped against a database comprising all available ASFLI-element sequences either from the *O. moubata* cell lines or from *O. porcinus* ticks using Bowtie2 (v.2.3.4.3) with default parameters. Mapped reads were assembled using SPAdes v3.11.1. Small RNA data were used with and without deduplication using BBMap version 38.38 (64). Reads were mapped against viral genomes or ASFLI-element-containing databases using Bowtie 2 (2.3.0) in Geneious with 22nt seed length for siRNA and 28nt seed length for piRNA and one allowed mismatch.

For the generation of full-length tick mitochondrial genomes, sequence data was mapped against a database containing all available *Ornithodoros* full-length mitochondrial sequences (as of 20 June 2019) using Newbler 3.0 and Bowtie 2 (2.3.0) and assembled using Newbler 3.0 and SPAdes v.3.11.1.

### Phylogenetic analysis

For identification of homologous genes from other viral genera, BLASTp (protein-protein BLAST) was used with amino acid sequences from the translated ASFV Ken06/Bus F1055L (helicase) and EP1242L (RNA polymerase subunit 2) genes. All viral sequences identified by BLASTp, covering at least 60% of the query, were used for further analysis. This yielded sequences from the NCLDV, including Faustoviruses, Pacmanvirus, Kaumoebavirus and Marseillevirus as well as African swine fever virus. These sequences together with the corresponding translation of the ASFLI-elements were aligned using MUSCLE in MEGA (65, 66). Nucleotide sequences of the EP1242L gene and equivalents from ASFV, Faustovirus, Pacmanvirus and Kaumoebavirus were also downloaded from GenBank. These nucleotide sequences together with the ASFLI-element were again aligned using MUSCLE in MEGA. A maximum likelihood tree was calculated from an alignment of NCLDV, ASFV and ASFV-like sequences in MEGA. The Jones-Taylor-Thornton (JTT) model allowing gamma distributed rates between sites (four categories), considering sites with >95 % data, and 100 bootstraps was used (67). More details about the methods used for the phylogenetic analyses are presented in Supplementary Appendix 1.

Tick mitochondrial genomes were aligned using MAFFT v7.388 in Geneious. The phylogenetic tree was constructed using IQ-TREE v1.6.5 with standard model selection, resulting in the best-fit model TIM2+F+R3 (AC=AT, CG=GT and unequal base frequencies + empirical base frequencies + FreeRate model with 3 categories).

### Clock rate estimates and Bayesian time scaled trees

To obtain an initial clock rate estimate, the ASFV nucleotide sequences were analysed using BEAST 1.8.4 (68, 69). Several different clock models were employed, and results were evaluated with the marginal likelihood estimation using path sampling and the stepping-stone sampling feature within BEAST (70). The clock models were: ‘strict’, ‘uncorrelated relaxed log-normal’, ‘uncorrelated relaxed gamma’, ‘uncorrelated relaxed exponential’, ‘random local clock’ and ‘fixed local clock’ (71–73). Apart from the clock models, the substitution model TN93 with gamma distributed site-to-site rate variation over four categories and constant population size tree prior model (effective population size) were used. Each BEAST run used a Markov-chain-Monte-Carlo (MCMC) chain with a length of 10,000,000 steps, sampling every 1,000 steps and discarding the first 10 % as burn-in.

All estimations of the time to the most recent common ancestor (TMRCA) are given in millions of years ago (mya). More details about the methods used for the molecular clock analyses are presented in Supplementary Appendix 1.

### Protein expression and purification in *E. coli* and rabbit immunisation

The reconstructed ASFLI-A104R gene (Supplementary Figure 2 N2/P2) was custom-synthesised and inserted into vector pMA-T (Geneart, Invitrogen). It was then amplified by PCR and the products were inserted into vector pMAL-p2X (New England Biolabs, Ipswich, Massachusetts). After transformation of *E. coli* (TB1) with the plasmid and induction with isopropyl-*β*-D-thiogalactopyranosid (IPTG), the maltose binding protein (MBP) fusion protein was purified by affinity solid phase extraction on an amylose resin as suggested by the manufacturer (New England Biolabs). Rabbits were immunised by intramuscular injections of 400 μg purified MBP–A104R fusion protein every four weeks over a period of four months. Blood was taken monthly and stored at 4 °C overnight after collection, followed by centrifugation for 10 min at 453 × *g* at room temperature. The resulting sera were stored at – 20 °C.

### Transfection

Approximately 10^6^ HEK 293T (ATCC® Number: CRL-11268™) cells were cultivated for 24 hours in 6-well plates. Using 5 μg of polyethylenimine (PEI) (74), the cells were transfected with 2.5 μg plasmid pCAGGS-Zecke-A104R in which transgene expression is directed by a hybrid HCMV major immediate early enhancer/chicken-actin promoter (75). The plasmid/PEI mix was added to the cell culture after pre-incubation at room temperature for 20 min and centrifuged at 600 × *g* for 60 min at 26 °C to enhance transfection efficacy. After 5 h of incubation at 37 °C, the medium was replaced by fresh medium, and after 48 h of further incubation, cells were harvested and lysed in 500 μl sample buffer for SDS-PAGE.

### SDS-PAGE and immunoblotting

SDS-PAGE was performed with a Mini-Protean Tetra System (Bio-Rad) using hand-cast SDS gels with 15 % polyacrylamide according to Laemmli (76). After separation, proteins were blotted onto nitrocellulose membranes (Whatman, Maidstone, United Kingdom) using a Trans-Blot SD Semi-Dry Transfer Cell (Bio-Rad) under conditions suggested by the manufacturer. Membranes were blocked by incubation in PBS supplemented with 6 % milk powder (Sigma Aldrich) and 10 % horse serum (Sigma Aldrich) overnight at 4 °C. After consecutive washes with 0.3% and 0.1% Tween20 in PBS, the blot was incubated with a 1:2,500 dilution of the rabbit serum (collected following the third immunisation), washed again with PBS-Tween20 as described above, and finally incubated with a 1:20,000 dilution of a peroxidase-conjugated secondary goat anti-rabbit antibody (Dianova, Hamburg, Germany). Signals were visualised using the ECL kit (Clarity Western ECL, Bio-Rad, Hercules, California) as suggested by the manufacturer.

### Statistical analysis

To calculate the number of ticks needed for the infection experiment, we conducted sample size calculation using R-studio (https://www.rstudio.com) as described elsewhere (77) with an assumed prevalence of 0.9 in one population and 0.3 in the second population. In order to obtain a power of 80 % (beta =0.2) and a confidence level of 99 % (alpha = 0.01), the required sample size was calculated to be ten animals per group.

## Declarations

### Ethics approval and consent to participate

All parts of the study involving animals were carried out following all applicable German animal welfare regulations and institutional guidelines. Immunisation of rabbits was reported to the competent authority under file accession LALLF-Nr. 7221.3-2.5-04/10.

### Consent for publication

Not applicable

### Availability of data and materials

ASFLI-element containing contigs and ASFV ken.re1 whole genome sequence are available from the International Nucleotide Sequence Database Collaboration (INSDC) databases under the study accession number PRJEB26739.

### Competing financial interest

The authors declare no competing financial interest

### Funding

JF was funded intramurally through the FLI ‘ASF-Research Network’ project. SL was founded by a BBSRC Roslin Institute Strategic Research program BBS/E/D/20002173 and University of Edinburgh Chancellor’s Fellowship. LBS was funded by UK BBSRC grant numbers BBS/E/1/00001741 and BB/P024270/1. AW and JK were funded by the Max Planck Society. This work was in part financed by COMPARE under the European Union’s Horizon 2020 programme, grant no. 643476, and by the German Federal Ministry of Education and Research-funded project DetektiVir, grant no. 13N13783.

### Authors’ contributions

JF, LF, HK, SB, DH and MB designed the experiments and did the comparative data analyses. JF conducted the qPCR, tick phylogeny and infection experiments and coordinated the study. JF, LF, SCS, AW, JK and DH conducted the NGS experiments and analysed the data. GMK did the cloning and expression of ASFV-like A104R. SL and JF did phylogeny and SL the BEAST analysis. LBS generated the tick cell lines and provided nucleic acid samples. JF, LF, SB and MB drafted the manuscript and LBS critically revised it. All authors commented on and approved the final manuscript.

## Acknowledgement

This work is dedicated to our friend and colleague Dr. Günther Keil who died on February 18^th^ 2020. The authors thank Carola Sauter-Louis (Friedrich-Loeffler-Institut, Germany) for help with the statistical analysis, Florian Pfaff (Friedrich-Loeffler-Institut, Germany) for the help with RNA sequencing, the Tick Cell Biobank for tick cell line provision, Ariane Düx (Robert-Koch-Institut, Germany) for help with the museum-stored field ticks, Guido Brandt and Cäcilia Freund (MPI-Jena, Germany) for their help with HiSeq-sequencing, Richard Bishop and Naftaly Githaka (ILRI-Kenya), Richard Lucius (Humboldt University Berlin-Germany), Fernando Boinas (University of Lisbon, Portugal), Sally Cutler (University of East London, UK), Cornelia Silaghi (Friedrich-Loeffler-Institut, Germany), Simon Carpenter (The Pirbright Institute, UK, under BBSRC project code BBS/E/I/00007039), Ricardo Péréz Sánchez (CSIC, Spain), Jason Dunlop (Berlin Museum for Natural History, Germany) and Laurence Vial (CIRAD, France) for supplying tick specimens and, furthermore, Juliane Horenk, Kathrin Giesow and Patrick Zitzow for technical assistance and René Klein for tick rearing.

