## Supplementary Material for "African swine fever virus-like integrated elements in a soft tick genome – an ancient virus vector arms race?"

† Dr. Günther Keil died on February 18th 2020

### Supplementary Figure 1

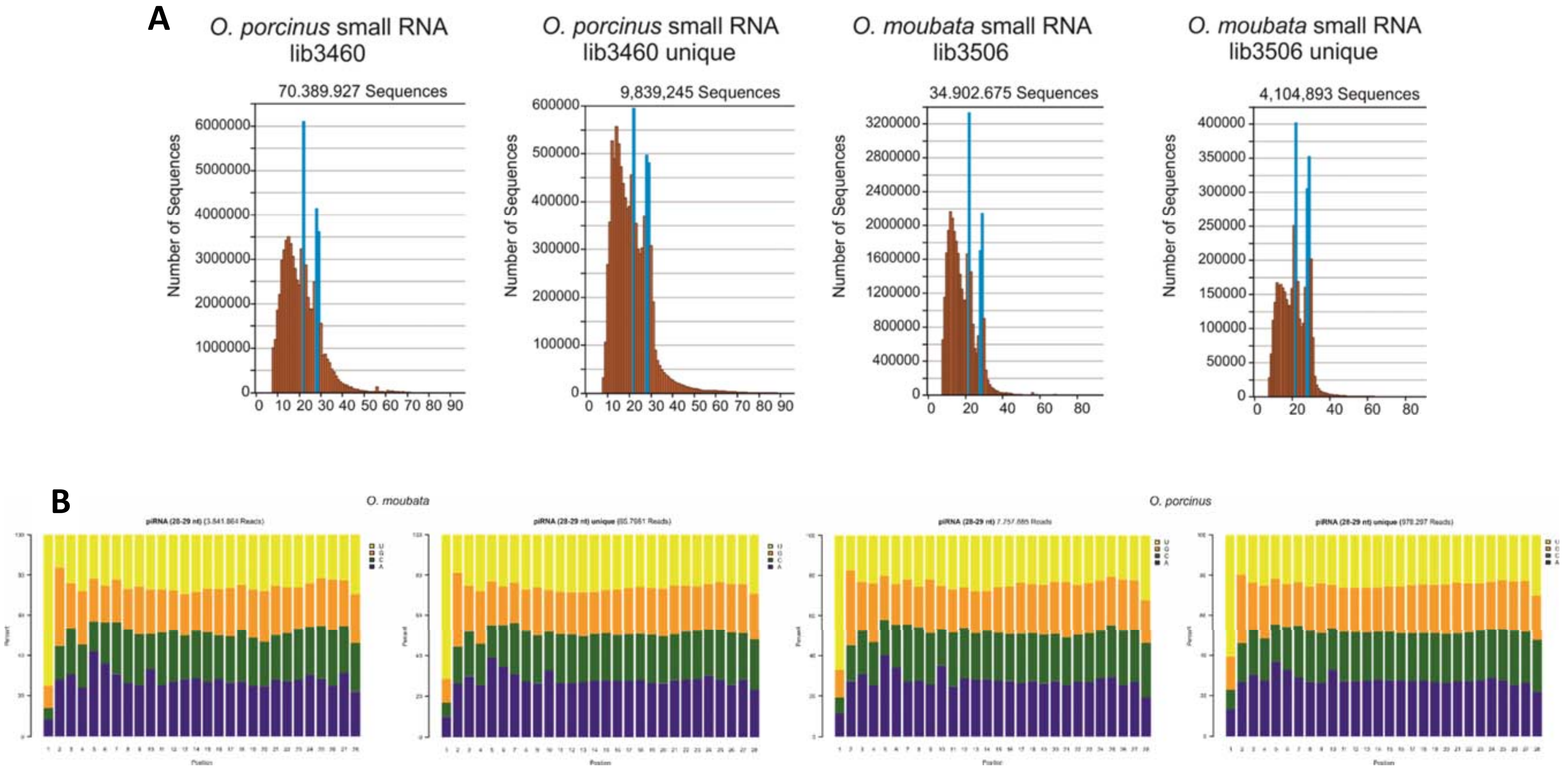

**Supplementary Figure 1 Read length distribution and piRNA nucleotide bias of small RNA libraries**
The length distribution is shown from small RNA libraries before and after removal of duplicate reads **(A)**. Reads of the piRNA fraction (28-29nt) were extracted from raw and deduplicated libraries and nucleotide frequencies for every position were calculated using R-studio (https://www.rstudio.com) **(B)**.

### Supplementary Figure 2

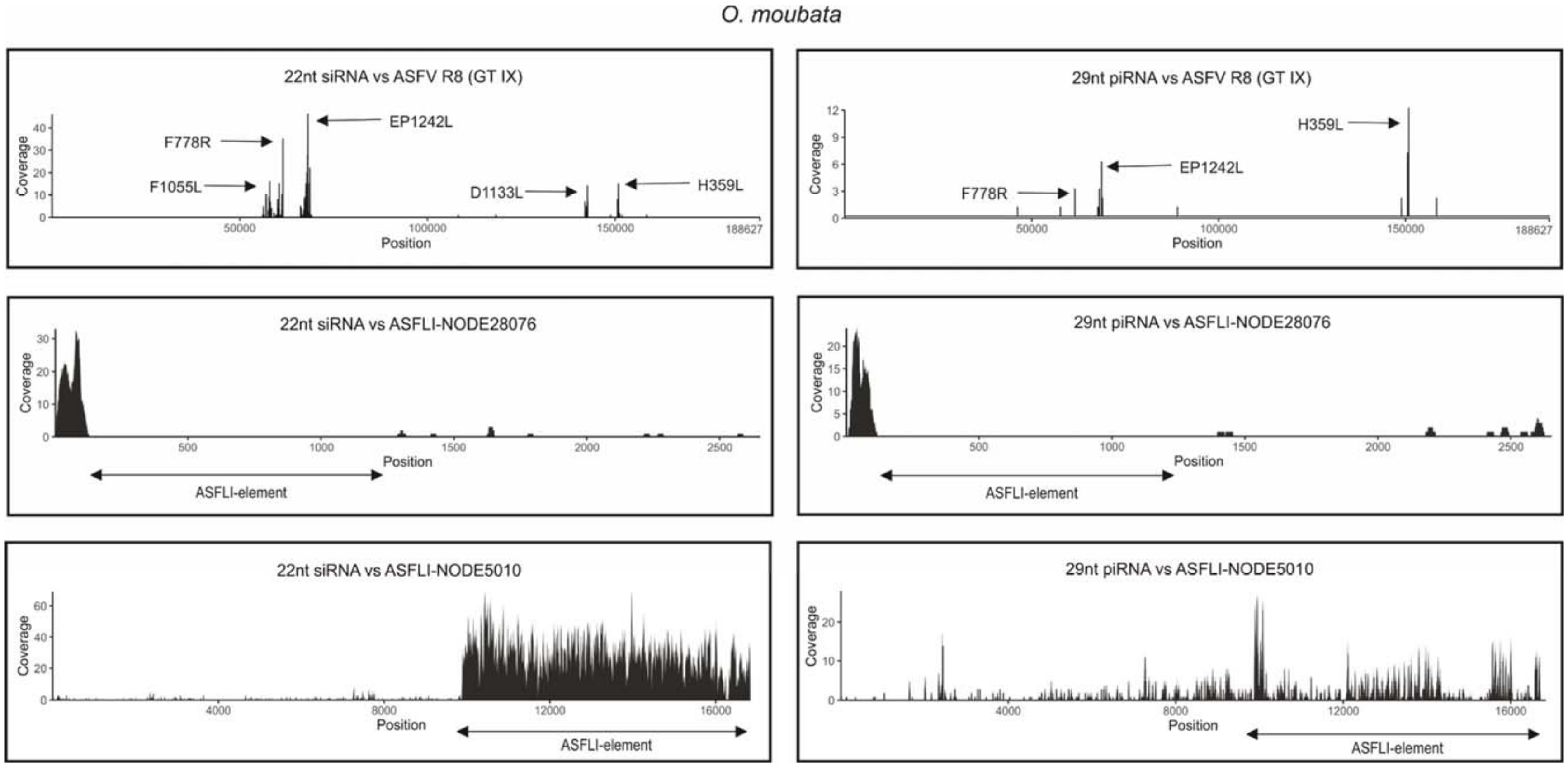

**Supplementary Figure 2. siRNA/piRNA mapping against ASFV-R8 (G IX) and two ASFLI-elements**

siRNA (22nt) and piRNA (29nt) fractions from *Ornithodoros moubata* were individually mapped against the ASFV-R8 (GT IX) whole-genome sequence and two *O. moubata* ASFLI-element-containing contigs NODE5010 and NODE28076 using Bowtie 2 (2.3.0) in Geneious. Marked (arrows) are the most abundant small RNA molecules related to the ASFV genes and ASFLI-elements.

### Supplementary Figure 3

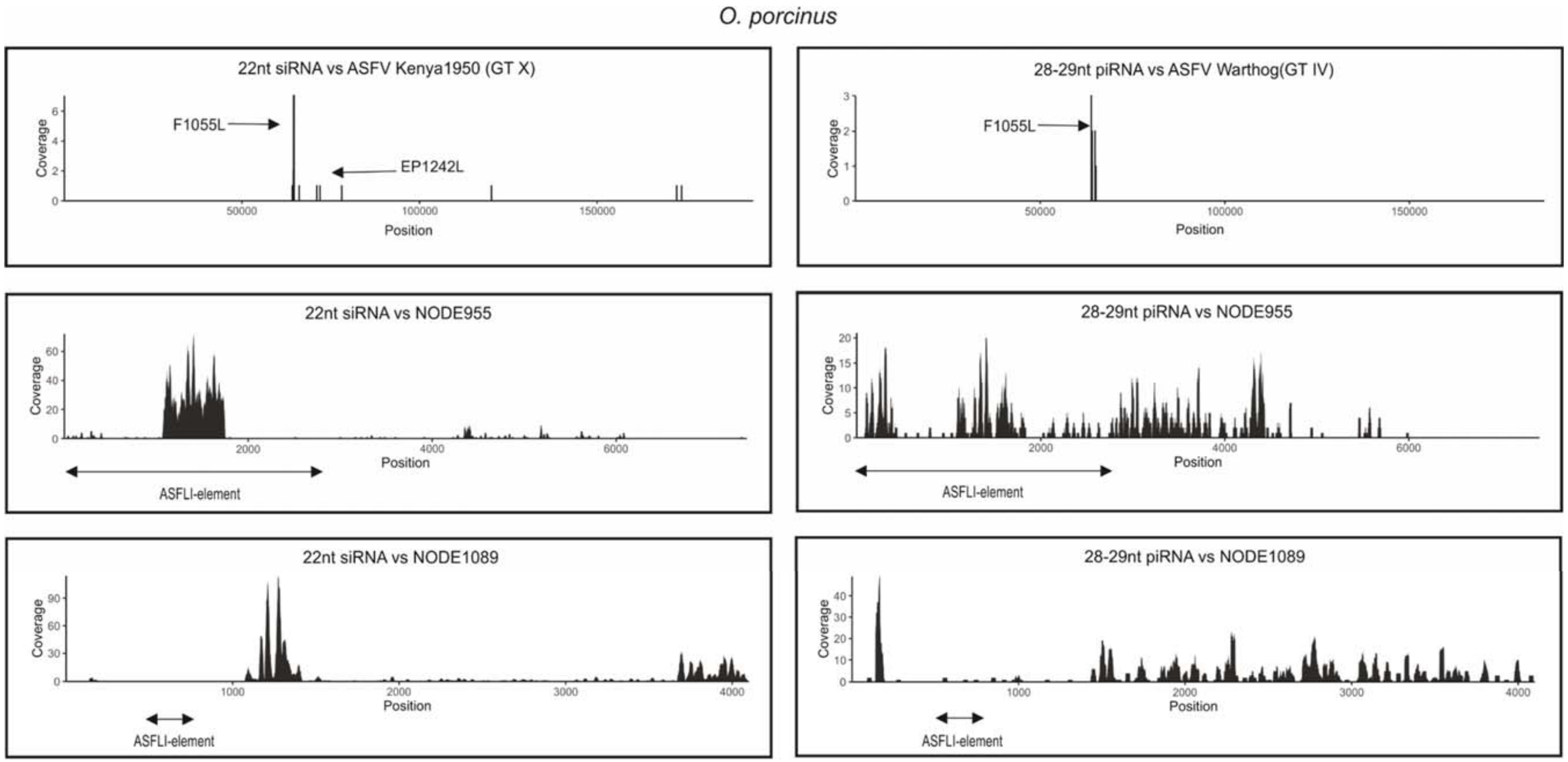

**Supplementary Figure 3. siRNA/piRNA mapping against ASFV-Kenya1950 (GT X), Warthog (G IV) and two ASFLI-elements**
siRNA (22nt) and piRNA (28-29nt) fractions from *Ornithodoros porcinus* were individually mapped against the ASFV-Kenya1950 (GT X), ASFV Wathog (GT IV) whole-genome sequence and two *O. porcinus* ASFLI-element containing contigs NODE955 and NODE1089 using Bowtie 2 (2.3.0) in Geneious. Marked (arrows) are the most abundant small RNA molecules relating to the ASFV genes and ASFLI-elements.

### Supplementary Figure 4

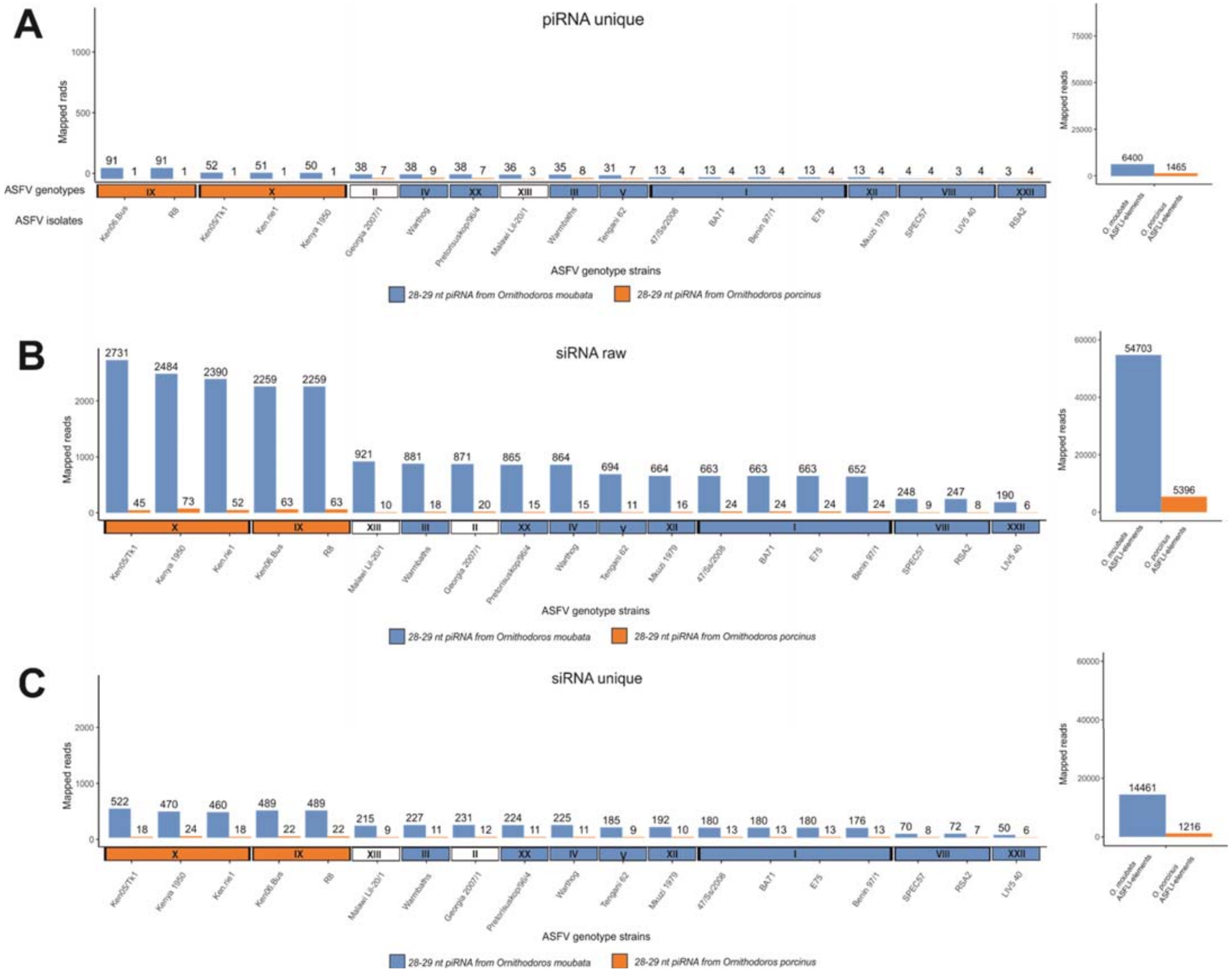

**Supplementary Figure 4. siRNA/piRNA mapping against ASFV and ASFLI-database**

siRNA (22nt) and piRNA (28-29nt) fractions from *Ornithodoros porcinus* and *Ornithodoros moubata*, before and after deduplication, were individually mapped against ASFV whole-genome sequence and the *O. moubata* or *O. porcinus* ASFLI-element containing datasets using Bowtie 2 (2.3.0) in Geneious.

### Supplementary Figure 5

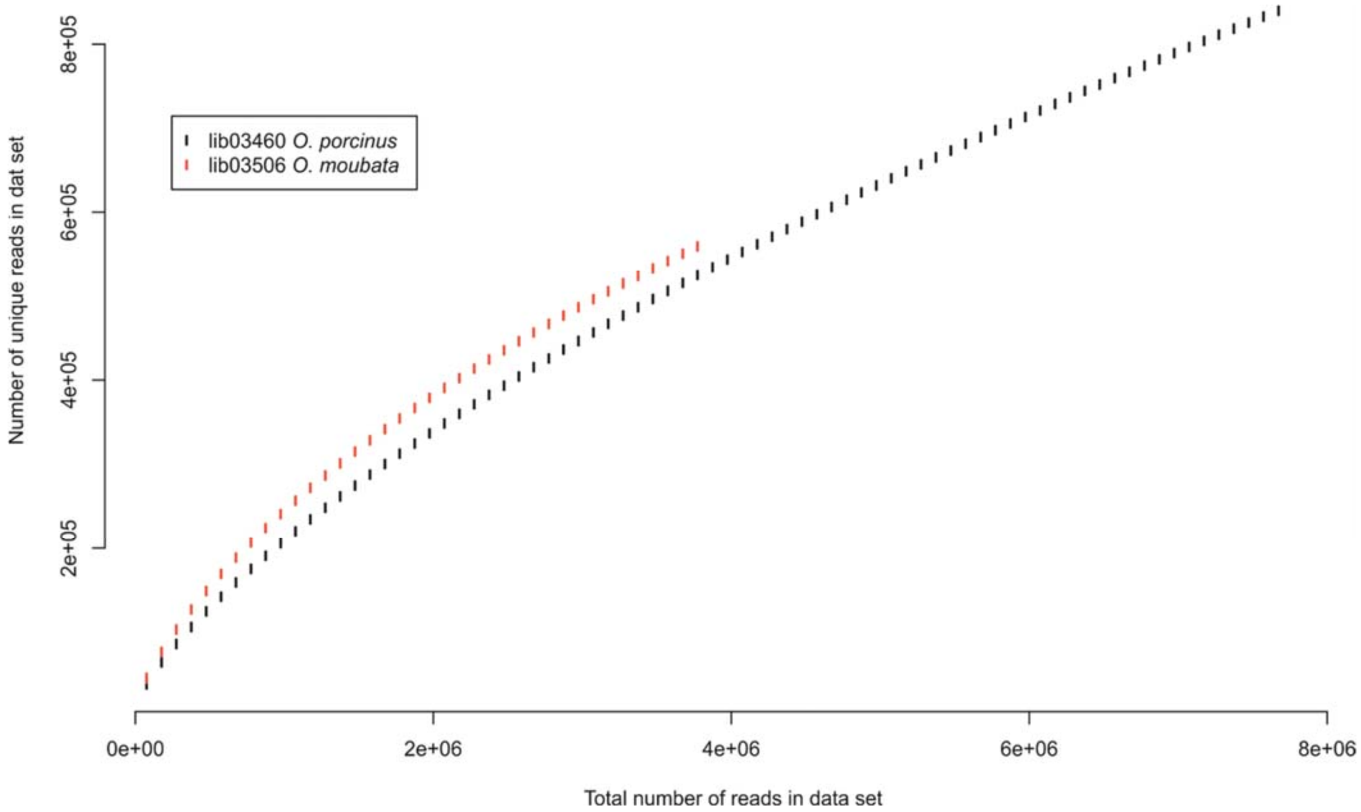

**Supplementary Figure 5. Rarefaction analysis of small RNA libraries lib03460 (*Ornithodoros porcinus*) and lib03506 (*Ornithodoros moubata*).**
To investigate the effect of amplification during library preparation both small RNA libraries were analysed and the results displayed using R-studio (https://www.rstudio.com).

### Supplementary Figure 6

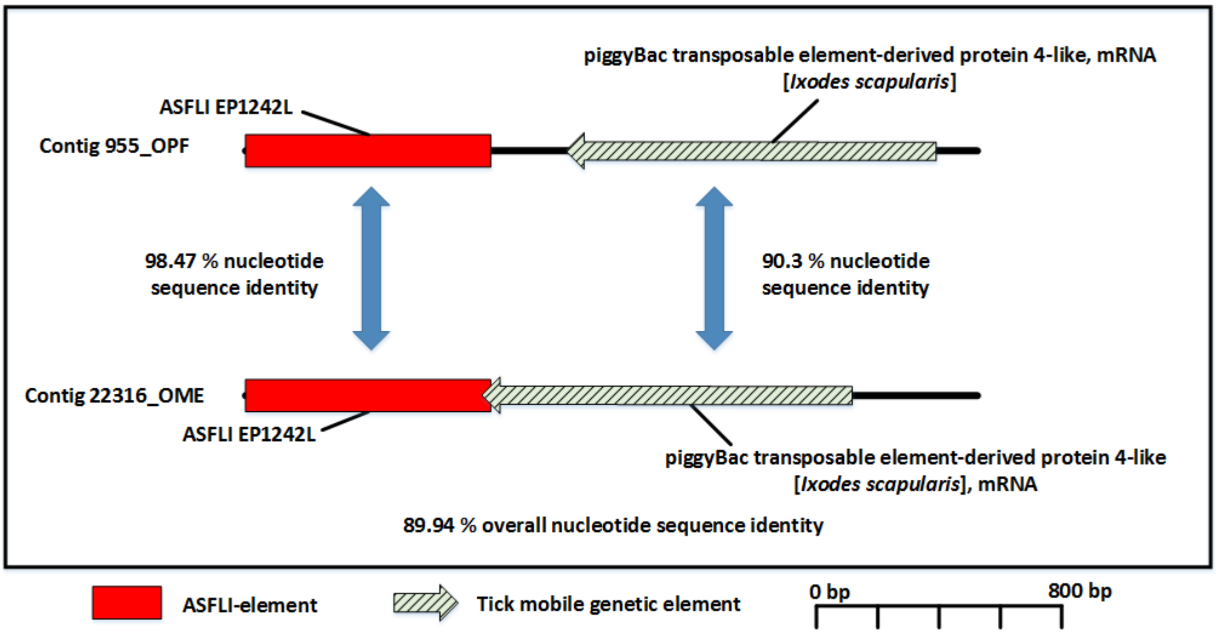

**Supplementary Figure 6. Similarity between ASFVLI contigs identified in *Ornithodoros moubata* cells and *Ornithodoros porcinus* ticks**
Contigs were assembled using SPAdes 3.13. on sequencing data obtained from *O. porcinus* ticks (OPF) and and OME/CTVM21 cells (OME )using default parameters. After contig identification by BLASTn and BLASTp search against the entire NCBI database and annotation, contigs were aligned using MAFFT v7.388 in Geneious.

### Supplementary Figure 7

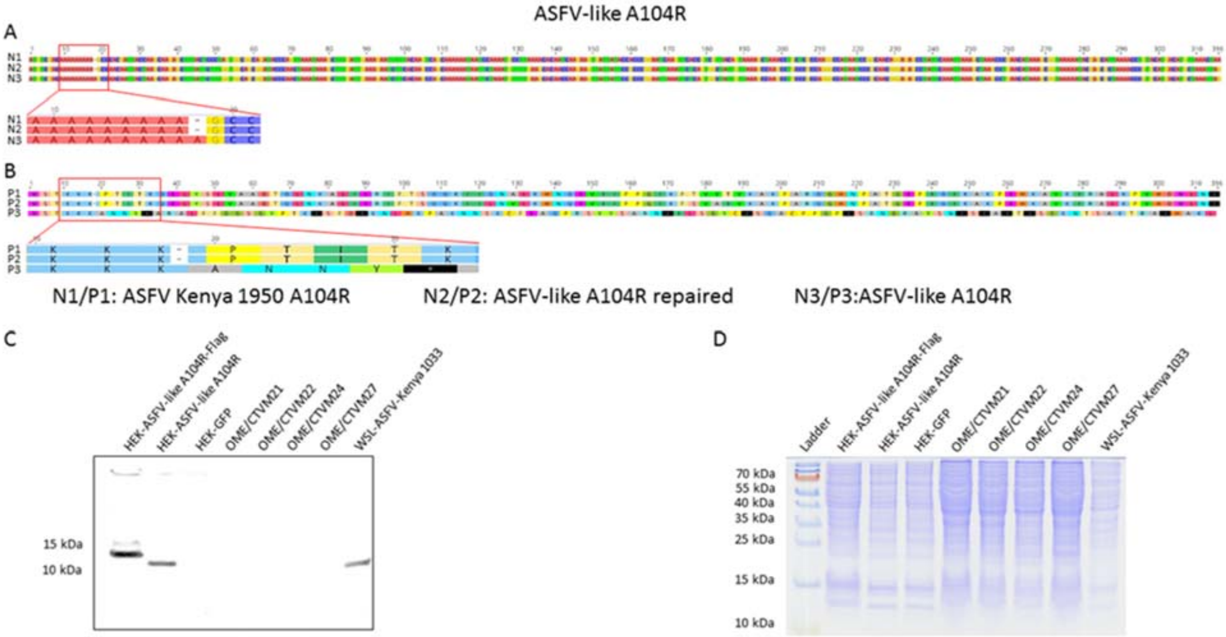

**Supplementary Figure 7. The reconstructed ASFV-like A104R protein is highly similar to its ASFV homologue**

**(A)** The ASFV-like A104R nucleotide sequence is highly similar to its ASFV-Kenya 1950 homologous A104R gene (83.0-85.0 %) with only one frameshift truncating the ORF. **(B)** When repaired, the ASFV-like A104R amino acid sequence is almost identical to its ASFV-Kenya 1950 A104R homologue (94.0-95.0 %). **(C)** A rabbit antiserum raised against the reconstructed A104 gene recognised a flag-tagged and an untagged version of A104R protein (lanes A104-Flag and A104, respectively), but showed no specific reaction with extracts of tick cell lines OME/CTVM21, OME/CTVM22, OME/CTVM24, and OME/CTVM27. In extracts of WSL-HP cells infected with ASFV Kenya 1033, the serum reacted with a single band of 12 kDa which is similar to the calculated molecular weight of ASFV A104R (11.6 kDa). **(D)** The Coomassie stained gel confirms equal loading with proteins.

### Supplementary Table 1

Supplementary Table 1

| Sample | Library | Type | NGS-Platform | Mode | Read length (bp) | Total reads | ASFV-like reads | % Identity to ASFV | Analysis | Reference |
|---|---|---|---|---|---|---|---|---|---|---|
| OME21 | Lib01543 | RNA | IonTorrent PGM | single end | 200-450 | 466.707 | 4 | 77.48-89.63 | Metagenome | This study |
| OME22 | Lib01544 | RNA | IonTorrent PGM | single end | 200-450 | 448.306 | 2 | 82.54-83.67 | Metagenome | This study |
| OME24 | Lib01545 | RNA | IonTorrent PGM | single end | 200-450 | 446.909 | 1 | 75.41 | Metagenome | This study |
| OME25 | Lib01610 | RNA | IonTorrent PGM | single end | 200-450 | 1.078.514 | 6 | 74.09-88.89 | Metagenome | This study |
| OME26 | Lib01611 | RNA | IonTorrent PGM | single end | 200-450 | 1.024.109 | 10 | 77.42-86.90 | Metagenome | This study |
| OME27 | Lib01546 | RNA | IonTorrent PGM | single end | 200-450 | 456.838 | 2 | 77.96-85.25 | Metagenome | This study |
| OME21 | Lib02151 | DNA | Illumina MiSeq | paired end | 300 | 6.966.615 | 1389 | 63.43-92.17 | Metagenome and assembly | This study |
| OME22 | Lib02152 | DNA | Illumina MiSeq | paired end | 300 | 7.967.586 | 1569 | 64.75-95.35 | Metagenome and assembly | This study |
| OME21 | Lib02339 | DNA | Illumina HiSeq | paired end | 50 | 277.000.000 | nd | nd | Assembly | This study |
| OME21 | LibMinION_1 | DNA | MinION | single molecule | ultra-long | 505.443 | nd | nd | Assembly | This study |
| OME21 | Lib02965 | mRNA | IonTorrent S5 | single end | 200 | 30.828.357 | 879 | 69.3 – 93.3 | Mapping | This study |
| OME21 | Lib02966/67 | Small RNA | IonTorrent S5 | single end | 200 | 59.966.291 | nd | nd | Mapping | This Study |
| O.porc | Lib03460 | Small RNA | IonTorrent S5 | single end | 200 | 70.389.927 | nd | nd | Mapping | This Study |
| O.moub | Lib03506 | Small RNA | IonTorrent S6 | single end | 200 | 34.902.675 | nd | nd | Mapping | This Study |
| O. moub. | N/A | midgut mRNA | Illumina HiSeq | paired end | 125 | 64.615.735 | 1593 | 71.00-91.20 | Mapping | Oleaga et al., 2017 |
| O. moub. | N/A | midgut mRNA | Illumina Genome Analyzer IIx | single end | 36 | 68.900.000 | 114 | nd | Mapping | Ogihara et al., 2015 |
| O.porc ken17 | Lib03101 | DNA | Illumina MiSeq | paired end | 300 | 7540296 | 227 | nd | Assembly and blast | This study |
| O.porc Fra19 | Lib03102 | DNA | Illumina MiSeq | paired end | 300 | 8506520 | 156 | nd | Assembly and blast | This study |
| Ornithodoros Mfn | Lib03076/GEA001 | DNA | Illumina HiSeq | paired end | 75 | 5978314 | 30 | nd | Mapping | This study |
| Ornithodoros Mfn | Lib03077/GEA002 | DNA | Illumina HiSeq | paired end | 75 | 8038160 | 21 | nd | Mapping | This study |
| Ornithodoros Mfn | Lib03078/MPU001.A01 | DNA | Illumina HiSeq | paired end | 75 | 5427474 | 12 | nd | Mapping | This study |
| Ornithodoros Mfn | Lib03079/NAL001.A01 | DNA | Illumina HiSeq | paired end | 75 | 7047107 | 0 | nd | Mapping | This study |
| Ornithodoros Mfn | Lib03080/KIS001.A01 | DNA | Illumina HiSeq | paired end | 75 | 7910598 | 18 | nd | Mapping | This study |
| Ornithodoros Mfn | Lib03081/AGL001.A01 | DNA | Illumina HiSeq | paired end | 75 | 5530990 | 136 | nd | Mapping | This study |
| Ornithodoros Mfn | Lib03082/TAB001.A01 | DNA | Illumina HiSeq | paired end | 75 | 5921628 | 3 | nd | Mapping | This study |
| Ornithodoros Mfn | Lib03083/TAB001.B01 | DNA | Illumina HiSeq | paired end | 75 | 8041845 | 3 | nd | Mapping | This study |
| Ornithodoros Mfn | Lib03087/GEA003.A01 | DNA | Illumina HiSeq | paired end | 75 | 9311389 | 6 | nd | Mapping | This study |
| Ornithodoros Mfn | Lib03084/MPA001.A01 | DNA | Illumina HiSeq | paired end | 75 | 5985437 | 32 | nd | Mapping | This study |
| Ornithodoros Mfn | Lib03085/MPA001.B01 | DNA | Illumina HiSeq | paired end | 75 | 5919694 | 25 | nd | Mapping | This study |
| Ornithodoros Mfn | Lib03086/NIA001.A01 | DNA | Illumina HiSeq | paired end | 75 | 7656401 | 44 | nd | Mapping | This study |
| Ornithodoros Mfn | AGL001.A0101 | DNA | Illumina HiSeq | paired end | 75 | 400.000.000 | nd | nd | Mapping, assembly and blast | This study |
| Ornithodoros Mfn | MPA001.A0101 | DNA | Illumina HiSeq | paired end | 75 | 400.000.000 | nd | nd | Mapping, assembly and blast | This study |

**Supplementary Table 1. Overview of NGS experiments conducted in this study and data obtained from the literature. *Ornithodoros moubata* cell line names have been abbreviated by removal of "/CTVM" and *Ornithodoros porcinus* and *Ornithodoros moubata* have been abbreviated for convenience.**

### Supplementary Table 2

| | | | | ASFLI-transcripts | | | | | | | |
|---|---|---|---|---|---|---|---|---|---|---|---|
| | Contigs | | | | blastn | | | | | | |
| Designation | Sample | Length | Coverage | Organism | ASFV-gene | Hit start | Hit end | Pairwise identity | E-value | Accession number | Data-reference |
| TK-Oleaga-1 | | 102 | | African swine fever virus strain Ken05/Tk1, complete genome | EP1242L | 69856 | 69957 | 91.2 % | 1.60e-30 | KM111294 | |
| TK-Oleaga-2 | | 146 | | African swine fever virus isolate Warmbaths, complete genome | MGF 360 15R | 49515 | 49584 | 82.9 % | 2.38e-09 | AY261365 | |
| TK-Oleaga-3 | | 177 | | African swine fever virus strain Ken06.Bus, complete genome | MGF 360 15R | 46341 | 46508 | 73.8 % | 4.02e-19 | KM111295 | |
| TK-Oleaga-5 | | 125 | | African swine fever virus isolate Kenya 1950, complete genome | EP1242L | 73220 | 73357 | 71.0 % | 5.59e-11 | AY261360 | |
| TK-Oleaga-10 | | 126 | | African swine fever virus isolate Malawi Lil-20/1 (1983), complete genome | F778R | 55696 | 55843 | 75.7 % | 1.83e-19 | AY261361 | |
| TK-Oleaga-11 | | 253 | | African swine fever virus isolate Kenya 1950, complete genome | F778R | 61516 | 61631 | 87.1 % | 8.88e-30 | AY261360 | |
| TK-Oleaga-12 | | 126 | | African swine fever virus strain Ken06.Bus, complete genome | F778R | 55936 | 56343 | 79.7 % | 4.29e-97 | KM111295 | |
| TK-Oleaga-13 | | 124 | | African swine fever virus isolate Kenya 1950, complete genome | F778R | 62373 | 62573 | 85.6 % | 6.38e-57 | AY261360 | |
| TK-Oleaga-14 | | 202 | | African swine fever virus isolate Warthog, complete genome | F778R | 55320 | 55471 | 79.3 % | 1.08e-09 | AY261366 | |
| TK-Oleaga-15 | O.moub | 395 | nd | African swine fever virus strain Ken06.Bus, complete genome | F165R | 55936 | 56343 | 79.7 % | 4.29e-97 | KM111295 | Oleaga et al. 2017 |
| TK-Oleaga-16 | | 1078 | | African swine fever virus isolate Kenya 1950, complete genome | F1055L | 63413 | 64540 | 76.0 % | 0 | AY261360 | |
| TK-Oleaga-17 | | 125 | | African swine fever virus strain Ken05/Tk1, complete genome | F1055L | 62073 | 62202 | 77.7 % | 6.40e-19 | KM111294 | |
| TK-Oleaga-18 | | 126 | | African swine fever virus isolate Kenya 1950, complete genome | F1055L | 64833 | 64943 | 85.6 % | 5.61e-26 | AY261360 | |
| TK-Oleaga-19 | | 125 | | African swine fever virus isolate Tengani 62, complete genome | F1055L | 57072 | 57202 | 79.4 % | 1.01e-22 | AY261364 | |
| TK-Oleaga-20 | | 126 | | African swine fever virus isolate Kenya 1950, complete genome | F1055L | 65698 | 65795 | 77.8 % | 2.55e-11 | AY261360 | |
| TK-Oleaga-21 | | 273 | | African swine fever virus isolate Malawi Lil-20/1 (1983), complete genome | F1055L | 60582 | 60884 | 72.6 % | 5.25e-39 | AY261361 | |
| TK-Oleaga-22 | | 176 | | African swine fever virus isolate Warmbaths, complete genome | E1242L | 67417 | 67598 | 84.1 % | 3.77e-47 | AY261365 | |
| TK-Oleaga-25 | | 295 | | African swine fever virus Georgia 2007/1 complete genome | H359L | 151371 | 151665 | 81.7 % | 2.71e-74 | FR682468 | |
| TK-Oleaga-26 | | 314 | | African swine fever virus isolate Warthog, complete genome | H359L | 148451 | 148759 | 79.3 % | 2.54e-68 | AY261366 | |
| TK-01610-R1 | | 233 | | African swine fever virus isolate Tengani 62, complete genome | B623R | 104199 | 103981 | 74.1 % | 3.00e-23 | AY261364 | |
| TK-01610-R2 | | 211 | | African swine fever virus isolate Kenya 1950, complete genome | P1192R | 153842 | 153976 | 88.9 % | 5.42e-39 | AY261360 | |
| TK-01610-R3 | OME25 | 202 | | African swine fever virus isolate Warthog, complete genome | H359L | 148576 | 148777 | 79.7 % | 3.66e-41 | AY261366 | |
| TK-01610-R4 | | 357 | | African swine fever virus isolate Pretorisuskop/96/4, complete genome | G1340L | 111013 | 110778 | 75.5 % | 4.17e-34 | AY261363 | |
| TK-01610-R5 | | 311 | | African swine fever virus isolate Warthog, complete genome | EP1242L | 66561 | 66445 | 78.6 % | 2.81e-17 | AY261366 | |
| TK-01610-R6 | | 286 | | African swine fever virus isolate Kenya 1950, complete genome | H124R | 157131 | 157077 | 81.8 % | 1.05e-03 | AY261360 | |
| TK-01611-R1 | | 323 | | African swine fever virus isolate Malawi Lil-20/1 (1983), complete genome | N/A | 184359 | 184575 | 77.4 % | 1.89e-38 | AY261361 | |
| TK-01611-R2 | | 346 | | African swine fever virus isolate Pretorisuskop/96/4, complete genome | H359L | 152815 | 153158 | 80.5 % | 3.51e-83 | AY261363 | |
| TK-01611-R3 | | 309 | | African swine fever virus isolate Kenya 1950, complete genome | Intergenic | 190031 | 189948 | 86.9 % | 4.90e-18 | AY261360 | |
| TK-01611-R4 | | 255 | | African swine fever virus isolate Kenya 1950, complete genome | B175L/B623R | 112128 | 111918 | 79.1 % | 9.44e-40 | AY261360 | |
| TK-01611-R5 | OME26 | 268 | | African swine fever virus isolate Warthog, complete genome | P1192R | 146033 | 145861 | 78.0 % | 4.58e-31 | AY261366 | |
| TK-01611-R6 | | 157 | | African swine fever virus isolate Malawi Lil-20/1 (1983), complete genome | MGF 360 15R | 47167 | 47233 | 82.6 % | 1.23e-06 | AY261361 | |
| TK-01611-R7 | | 83 | 1 | African swine fever virus isolate Pretorisuskop/96/4, complete genome | MGF 360 15R | 50065 | 50114 | 84.3 % | 2.22e-03 | AY261363 | This study |
| TK-01611-R8 | | 316 | | African swine fever virus isolate Kenya 1950, complete genome | F778R | 61156 | 61004 | 86.3 % | 6.36e-42 | AY261360 | |
| TK-01611-R9 | | 318 | | African swine fever virus isolate Pretorisuskop/96/4, complete genome | H359L | 153157 | 152826 | 77.7 % | 1.60e-68 | AY261363 | |
| TK-01611-R10 | | 230 | | African swine fever virus isolate Warthog, complete genome | H359L | 148091 | 148317 | 80.7 % | 1.31e-50 | AY261366 | |
| TK-01544-46-R1 | | 212 | | African swine fever virus isolate Pretorisuskop/96/4, complete genome | H359L | 152916 | 153062 | 83.7 % | 7.26e-53 | AY261363 | |
| TK-01544-46-R2 | OME22,24,27 | 336 | | African swine fever virus Benin 97/1 pathogenic isolate, complete genome | G1340L | 104523 | 104582 | 82.5 % | 5.26e-05 | AM712239 | |
| TK-01544-46-R3 | | 271 | | African swine fever virus strain Ken05/Tk1, complete genome | F1055L | 62437 | 62678 | 75.4 % | 1.15e-38 | KM111294 | |
| TK-01544-46-R4 | | 296 | | African swine fever virus isolate Kenya 1950, complete genome | F1055L | 63996 | 64308 | 78.0 % | 8.27e-66 | AY261360 | |
| TK-01544-46-R5 | | 249 | | African swine fever virus strain Ken06.Bus, complete genome | H359L | 149108 | 149168 | 85.2 % | 2.89e-08 | KM111295 | |
| TK-01543-R1 | | 274 | | African swine fever virus isolate Warmbaths, complete genome | MGF 360 15R | 49515 | 49622 | 77.8 % | 1.77e-13 | AY261365 | |
| TK-01543-R2 | OME21 | 307 | | African swine fever virus isolate Warmbaths, complete genome | EP1242L | 67601 | 67407 | 83.2 % | 1.36e-46 | AY261365 | |
| TK-01543-R3 | | 407 | | African swine fever virus Georgia 2007/1 complete genome | P1192R | 149403 | 149566 | 89.6 % | 4.17e-53 | FR682468 | |
| TK-01543-R4 | | 208 | | African swine fever virus Georgia 2007/1 complete genome | H171R | 152492 | 152388 | 77.5 % | 9.19e-11 | FR682468 | |

| | | | | | | | | | | | |
|---|---|---|---|---|---|---|---|---|---|---|---|
| TK_02965_1 | | 113 | | African swine fever virus strain R35, complete genome | EP1242L | 67232 | 67344 | 90.27 % | 5,00E-34 | MH025920.1 | |
| TK_02965_2 | | 143 | | African swine fever virus strain R35, complete genome | EP1242L | 67571 | 67712 | 89.51% | 1,00E-43 | MH025920.1 | |
| TK_02965_3 | | 120 | | African swine fever virus strain Ken05/Tk1, complete genome | EP1242L | 69827 | 69946 | 90.00% | 1,00E-36 | KM111294.1 | |
| TK_02965_4 | | 136 | | African swine fever virus strain Ken05/Tk1, complete genome | EP1242L | 70088 | 70223 | 93.38% | 4,00E-49 | KM111294.1 | |
| TK_02965_5 | | 142 | | African swine fever virus isolate Pretorisuskop/96/4, complete genome | EP84R | 71152 | 72194 | 90.2 % | 2.47e-45 | AY261363 | |
| TK_02965_6 | | 139 | | African swine fever virus isolate Malawi Lil-20/1 (1983), complete genome | EP84R | 68263 | 68401 | 90.6 % | 2.40e-45 | AY261361 | |
| TK_02965_7 | | 170 | | African swine fever virus Benin 97/1 | MGF 360 15R | 43917 | 43986 | 84.3 % | 1.73e-10 | AM712239 | |
| TK_02965_8 | | 167 | | African swine fever virus strain Ken06.Bus, complete genome | MGF 360 15R | 46341 | 46492 | 73.7 % | 6.3e-16 | KM111295 | |
| TK_02965_9 | | 236 | | African swine fever virus isolate Tengani 62, complete genome | MGF 360 15R | 45263 | 45360 | 69.3 % | 3.59e-2 | AY261364 | |
| TK_02965_10 | OME21 | 636 | n.d. | African swine fever virus strain Ken06.Bus, complete genome | EP1242L | 66267 | 66596 | 77.9 % | 4.25e-70 | KM111295 | This study |
| TK_02965_11 | | 170 | | African swine fever virus isolate Kenya 1950, complete genome | F778R | 61521 | 61706 | 79.6 % | 4.34e-37 | AY261360 | |
| TK_02965_12 | | 169 | | African swine fever virus isolate Kenya 1950, complete genome | F778R | 61865 | 61981 | 80.3 % | 8.32e-23 | AY261360 | |
| TK_02965_13 | | 274 | | African swine fever virus strain Ken05/Tk1, complete genome | F778R | 59523 | 59794 | 85.4 % | 9.95e-80 | KM111294 | |
| TK_02965_14 | | 679 | | African swine fever virus isolate Kenya 1950, complete genome | F778R/F165R | 62400 | 63094 | 81.1 % | 0 | AY261360 | |
| TK_02965_15 | | 1193 | | African swine fever virus strain BA71V, complete genome | F1055L | 42983 | 44116 | 75.5 % | 0 | ASU18466 | |
| TK_02965_16 | | 1697 | | African swine fever virus isolate Kenya 1950, complete genome | F1055L | 64534 | 66227 | 70.0 % | 0 | AY261360 | |
| TK_02965_17 | | 134 | | African swine fever virus strain Ken05/Tk1, complete genome | B263R | 109618 | 109503 | 75.00% | 4,00E-11 | KM111294.1 | |
| TK_02965_18 | | 117 | | African swine fever virus strain L60, complete genome | | 103252 | 103164 | 76.40% | 2,00E-07 | KM262844.1 | |
| TK_02965_19 | | 256 | | African swine fever virus isolate Malawi Lil-20/1 (1983), complete genome | P1192R | 158148 | 158037 | 91.96% | 9,00E-36 | AY261361.1 | |
| TK_02965_20 | | 373 | | African swine fever virus isolate Tengani 62, complete genome | H359L | 146881 | 147255 | 82.13% | 2,00E-97 | AY261364.1 | |

**Supplementary Table 2. ASFV-like transcripts detected in data from six *Ornithodoros moubata* cell lines and *O.moubata* ticks by mRNA-sequencing and data analysis from published sequences**
ASFV-like sequences from tick cell libraries lib01543-46 und lib01610-11 (total RNA) were identified using RIEMS. The resulting reads were blasted against the NCBI BLASTn database for confirmation and annotation. *Ornithodoros moubata* transcriptome data from a previous study[42] was downloaded from Genbank, and ASFV-like sequences were identified by mapping the data against a tailored database containing all known ASFV sequences and all previously obtained ASFV-like sequences. The resulting reads and contigs were annotated after a BLASTn search against the entire BLASTn database. Sequencing data from the mRNA enriched OME21 library lib02965 was mapped against a tailored database containing all known ASFV sequences and all previously obtained ASFV-like sequences, and resulting contigs were blasted against the entire BLASTn database. Per sequence, only the hit with the lowest e-value is shown.

### Supplementary Table 3

| SPAdes contigs with ASFLI-elements (>500bp) | | | | | | | | | |
|---|---|---|---|---|---|---|---|---|---|
| Contigs | | | blastn-results | | | | | | |
| Designation | Length | Coverage | Organism/Gene | ASFV gene | Query start | Query end | Pairwise identity | E-value | Accession number |
| Contig_885 | 40590 | 7.4 | African swine fever virus isolate Malawi Lil-20/1 (1983), complete genome | CP312R-O174L-O61R | 15673 | 17203 | 69.9% | 0 | AY261361 |
| | | | African swine fever virus isolate Malawi Lil-20/1 (1983), complete genome | CP530R-CP80R-CP312R | 17456 | 17966 | 70.0% | 1.64e-51 | AY261361 |
| | | | Ixodes scapularis BAC ISG1-01F14, complete sequence | / | 27406 | 27636 | 76.6% | 3.87e-34 | AC205630 |
| | | | Ixodes scapularis, clone XX-43A1, complete sequence | / | 29281 | 29416 | 71.1% | 6.56e-06 | AC192419 |
| | | | Ixodes scapularis, clone XX-43A1, complete sequence | / | 30865 | 31116 | 71.1% | 1.27e-08 | AC192419 |
| | | | Ixodes scapularis conserved hypothetical protein, mRNA | / | 35380 | 35569 | 72.6% | 7.48e-18 | XM_002411761 |
| Contig_5010 | 16841 | 16.2 | Ixodes scapularis clone 01_G21_T3 tandem repeat region | | 111 | 189 | 79.7% | 7.78e-07 | GU318717 |
| | | | African swine fever virus strain Ken05/Tk1, complete genome | EP1242L-EP84R-EP424R | 9319 | 12322 | 79.4% | 0 | KM111294 |
| | | | African swine fever virus isolate Kenya 1950, complete genome | F778R-F165R-F1055L | 12322 | 14945 | 73.8% | 0 | AY261360 |
| Contig_11452 | 8635 | 4.9 | African swine fever virus isolate Malawi Lil-20/1 (1983), complete genome | CP312R-O174L | 369 | 1031 | 69.3% | 5.51e-75 | AY261361 |
| | | | African swine fever virus isolate Tengani 62, complete genome | CP312R | 1166 | 1332 | 82.6% | 5.52e-37 | AY261364 |
| | | | African swine fever virus isolate Malawi Lil-20/1 (1983), complete genome | CP530R-CP80R-CP312R | 1576 | 2028 | 70.6% | 2.06e-42 | AY261361 |
| | | | African swine fever virus isolate Malawi Lil-20/1 (1983), complete genome | B263R | 3143 | 3385 | 72.5% | 1.39e-25 | AY261361 |
| | | | Ixodes scapularis PrC protein, putative, mRNA | / | 5633 | 5779 | 73.7% | 7.69e-10 | XM_002402644 |
| Contig_11845 | 8352 | 4.1 | African swine fever virus strain Ken05/Tk1, complete genome | A179L | 28 | 309 | 82.1% | 1.21e-70 | KM111294 |
| | | | African swine fever virus Georgia 2007/1 complete genome | A859L | 386 | 1464 | 76.7% | 0 | FR682468 |
| | | | African swine fever virus strain Ken05/Tk1, complete genome | A859L | 1237 | 1317 | 77.8% | 7.07e-06 | KM11294 |
| | | | African swine fever virus strain Ken06.Bus, complete genome | A238L-A859L | 1914 | 2680 | 78.2% | 0 | KM11295 |
| | | | African swine fever virus IkB-like protein Ug5ELp28 (5EL) gene | A238L-A859L | 2897 | 3467 | 73.1% | 3.97e-83 | AF014483 |
| | | | African swine fever virus strain L60, complete genome | MGF36015R | 3975 | 4726 | 67.8% | 4.84e-63 | KM262844 |
| Contig_12772 | 7718 | 13.4 | African swine fever virus strain Ken06.Bus, complete genome | D1133L | 839 | 2305 | 78.9% | 0 | KM111295 |
| | | | African swine fever virus strain Ken05/Tk1, complete genome | H171R-H24R | 2313 | 2714 | 65.0% | 1.69e-24 | KM111294 |
| | | | African swine fever virus isolate Pretorisuskop/96/4, complete genome | H359L | 2764 | 2982 | 80.08% | 7.67e-48 | AY261363 |
| | | | African swine fever virus Georgia 2007/1 complete genome | H171R | 2983 | 3238 | 79.5% | 1.48e-50 | FR682468 |
| | | | African swine fever virus strain Ken06.Bus, complete genome | H171R-H124R | 3265 | 3626 | 62.9% | 6.87e-10 | KM111295 |
| | | | African swine fever virus isolate Warthog, complete genome | H359L-H171R | 3632 | 4880 | 80.3% | 0 | AY261366 |
| | | | African swine fever virus strain Ken05/Tk1, complete genome | F778R-F165R | 7592 | 7716 | 84% | 1.14e-26 | KM111294 |
| Contig_15264 | 6370 | 4.0 | African swine fever virus isolate Pretorisuskop/96/4, complete genome | G1340L | 573 | 807 | 75.4% | 2.86e-34 | AY261363 |
| | | | African swine fever virus isolate Kenya 1950, complete genome | B175L-B263R-B66L-G1340L | 3167 | 4952 | 69.7% | 0 | AY261360 |
| | | | African swine fever virus isolate Kenya 1950, complete genome | C962R | 4956 | 5117 | 77.6% | 1.69e-24 | AY261360 |
| Contig_17249 | 5489 | 4.2 | African swine fever virus strain Ken06.Bus, complete genome | G1211R-CP123L | 3211 | 3485 | 76.1% | 3.97e-45 | KM111295 |
| | | | African swine fever virus isolate Warthog, complete genome | P1192R | 3686 | 3849 | 78.0% | 2.20e-29 | AY261366 |
| | | | African swine fever virus isolate Kenya 1950, complete genome | P1192R | 3855 | 4040 | 86.6% | 3.48e-52 | AY261360 |
| | | | African swine fever virus isolate Malawi Lil-20/1 (1983), complete genome | Q706L-QP509L | 4036 | 4308 | 84.7% | 2.50e-79 | AY261361 |
| | | | African swine fever virus isolate Warthog, complete genome | H359L | 4346 | 5298 | 80.3% | 0 | AY261366 |
| Contig_18126 | 5152 | 26.6 | African swine fever virus isolate Warthog, complete genome | A104R-A240L | 2981 | 3833 | 71.3% | 1.38e-120 | AY261366 |
| | | | African swine fever virus strain Ken06.Bus, complete genome | A224L | 4246 | 4732 | 68.1% | 9.98e-53 | KM111295 |
| | | | Ixodes scapularis BAC ISG1-01F14, complete sequence | / | 4867 | 5152 | 68.5% | 6.74e-17 | AC205630 |
| Contig_19570 | 4637 | 5.3 | African swine fever virus Georgia 2007/1 complete genome | CP530R-CP80R-CP312R | 149 | 724 | 68.8% | 3.48e-52 | FR682468 |
| | | | Ixodes scapularis BAC ISG1-15P02, complete sequence | / | 901 | 933 | 93.9% | 3.73e-01 | AC205638 |
| | | | African swine fever virus isolate Malawi Lil-20/1 (1983), complete genome | CP312R-O174L | 934 | 1963 | 69.3% | 5.15e-126 | AY261361 |
| | | | African swine fever virus BA71V protein p12 (O61R) gene, complete cds | O61R | 1858 | 1963 | 79.2% | 1.22e-13 | M84186 |

| | | | | | | | | | |
|---|---|---|---|---|---|---|---|---|---|
| Contig_21116 | 3824 | 4.2 | African swine fever virus isolate Pretorisuskop/96/4, complete genome | C962R | 233 | 1058 | 72.7% | 1.21e-127 | AY261363 |
| | | | African swine fever virus strain Ken06.Bus, complete genome | C962R | 1062 | 1749 | 78.0% | 3.04e-154 | KM111295 |
| | | | Ixodes scapularis BAC ISG1-42P02, complete sequence | / | 2585 | 3103 | 64.4% | 2.20e-10 | AC205646 |
| Contig_22316 | 3339 | 7.3 | African swine fever virus isolate Pretorisuskop/96/4, complete genome | EP1242L-EP84R-EP424R | 2632 | 3459 | 80.4% | 0 | AY261363 |
| | | | African swine fever virus isolate Warthog, complete genome | EP1242L | 3608 | 3823 | 80.8% | 4.24e-51 | AY261366 |
| Contig_24457 | 2653 | 7.9 | African swine fever virus isolate Warthog, complete genome | EP1242L | 1 | 676 | 76.9% | 1.38e-158 | AY261366 |
| Contig_28076 | 2653 | 7.9 | African swine fever virus isolate Pretorisuskop/96/4, complete genome | C147L-C62L-C962R | 123 | 763 | 72.0% | 2.33e-86 | AY261363 |
| | | | African swine fever virus Georgia 2007/1 complete genome | C315R | 769 | 865 | 79.4% | 5.50e-12 | FR682468 |
| | | | African swine fever virus strain Ken06.Bus, complete genome | C475L | 861 | 1242 | 76.9% | 7.13e-74 | KM111295 |
| Contig_29175 | 2475 | 6.8 | African swine fever virus isolate Tengani 62, complete genome | H359L | 4 | 675 | 82.7% | 0 | AY261364 |
| Contig_31135 | 2196 | 2.5 | African swine fever virus isolate Kenya 1950, complete genome | F778R | 2 | 447 | 84.3% | 5.84e-132 | AY261360 |
| | | | African swine fever virus strain Ken05/Tk1, complete genome | F778R-F165R | 476 | 2177 | 76.4% | 0 | KM111294 |
| | | | African swine fever virus LMW6DL origin binding protein gene, complete cds | F165R | 1904 | 2177 | 70.8% | 3.04e-34 | L12174 |
| Contig_35077 | 1763 | 23.9 | African swine fever virus isolate Kenya 1950, complete genome | F778R | 496 | 1040 | 84.0% | 3.68e-166 | AY261360 |

**SPAdes contigs with EASFL-elements (150-500bp)**

| Contigs | | | blastn-results | | | | | | |
|---|---|---|---|---|---|---|---|---|---|
| **Designation** | **Length** | **Coverage** | **Organism/Gene** | **ASFV gene** | **Query start** | **Query end** | **Pairwise identity** | **E-value** | **Accession number** |
| Contig_79 | 85582 | 6.2 | African swine fever virus OURT 88/3 (avirulent field isolate), complete genome | B602L | 72104 | 72270 | 69.5% | 3.98e-06 | AM712240 |
| | | | Ixodes scapularis hypothetical protein, mRNA | / | 82534 | 82592 | 91.5% | 7.68e-09 | XM_002406158 |
| | | | Ixodes scapularis clone 01_N16_T3 tandem repeat region | / | 83215 | 83297 | 75.9% | 7.19e-03 | GU319006 |
| | | | Ixodes scapularis clone 01_N16_T3 tandem repeat region | / | 51068 | 51149 | 77.1% | 7.19e-03 | GU319006 |
| Contig_1333 | 34743 | 6.0 | African swine fever virus isolate Malawi Lil-20/1 (1983), complete genome | B263R | 1 | 168 | 72.4% | 6.01e-12 | AY261361 |
| | | | Ixodes scapularis oligopeptide transporter, putative, mRNA | / | 2214 | 2289 | 85.5% | 2.10e-11 | XM_002414423 |
| | | | Ixodes scapularis oligopeptide transporter, putative, mRNA | / | 3122 | 3248 | 73.2% | 3.80e-08 | XM_002414423 |
| | | | Ixodes scapularis oligopeptide transporter, putative, mRNA | / | 13290 | 13393 | 76.0% | 1.32e-07 | XM_002414423 |
| Contig_5756 | 15286 | 5.3 | African swine fever virus isolate Warthog, complete genome | B263R | 5260 | 5595 | 71.3% | 1.20e-35 | AY261366 |
| Contig_6417 | 14138 | 5.8 | African swine fever virus isolate Warthog, complete genome | B263R | 8106 | 8495 | 71.0% | 2.78-34 | AY261366 |
| Contig_8979 | 10836 | 16.6 | African swine fever virus isolate Malawi Lil-20/1 (1983), complete genome | Q706L-QP509L | 4 | 197 | 84.6% | 4.71e-53 | AY261361 |
| Contig_10451 | 9433 | 18.9 | African swine fever virus isolate Warthog, complete genome | B263R | 1609 | 1999 | 71.3% | 2.29e-44 | AY261366 |
| Contig_11363 | 8706 | 7.2 | African swine fever virus Georgia 2007/1 complete genome | B263R | 559 | 805 | 71.8% | 9.72e-24 | FR682468 |
| | | | Ixodes scapularis clone 01_A18_T7 tandem repeat region | / | 5070 | 5325 | 68.2% | 7,00E-13 | GU318451 |
| | | | Ixodes scapularis, clone XX-5A1, complete sequence | / | 4955 | 5325 | 65.2% | 8.53e-12 | AC192414 |
| Contig_12106 | 8169 | 11.0 | African swine fever virus isolate Malawi Lil-20/1 (1983), complete genome | Q706L-QP509L | 7892 | 8166 | 84.4% | 8.30e-79 | AY261361 |
| | | | African swine fever virus isolate Kenya 1950, complete genome | P1192R | 7753 | 7896 | 89.6% | 1.61e-43 | AY261360 |
| Contig_13936 | 7045 | 7.1 | African swine fever virus isolate Malawi Lil-20/1 (1983), complete genome | B263R | 4157 | 4545 | 69.3% | 2.72e-34 | AY261361 |
| Contig_16516 | 5808 | 4.7 | African swine fever virus isolate Warthog, complete genome | B263R | 261 | 580 | 70.08% | 3.31e-33 | AY261366 |
| Contig_16776 | 5688 | 5.4 | African swine fever virus isolate Warthog, complete genome | B263R | 4410 | 4804 | 72.1% | 1.41e-50 | AY261366 |
| Contig_17474 | 5396 | 4.2 | African swine fever virus isolate Tengani 62, complete genome | MGF360-15R | 1354 | 1792 | 66.8% | 1.40e-29 | AY261364 |
| | | | African swine fever virus isolate Kenya 1950, complete genome | MGF360-15R | 1099 | 1245 | 74.1% | 1.40e-10 | AY261360 |
| Contig_18735 | 4920 | 9.0 | African swine fever virus isolate Malawi Lil-20/1 (1983), complete genome | CP530R-CP80R-CP312R | 2009 | 2461 | 70.3% | 3.76e-43 | AY261361 |
| Contig_27716 | 2712 | 2.7 | African swine fever virus strain Ken06.Bus, complete genome | EP1242L | 1 | 250 | 78.6% | 2.65e-56 | KM111295 |
| Contig_31254 | 2179 | 6.8 | African swine fever virus strain Ken06.Bus, complete genome | G1211R-CP123L | 214 | 461 | 77.8% | 6.67e-45 | KM111295 |
| | | | African swine fever virus isolate Warmbaths, complete genome | NP868R | 465 | 591 | 73.9% | 8.69e-12 | AY261365 |
| Contig_32100 | 2079 | 4.0 | African swine fever virus isolate Mkuzi 1979, complete genome | G1340L | 6 | 372 | 72.6% | 2.33e-44 | AY261362 |
| | | | Ixodes scapularis conserved hypothetical protein, mRNA | / | 1143 | 1223 | 78.0% | 3.46e-04 | XM_002399220 |
| Contig_41411 | 971 | 9.6 | African swine fever virus isolate Warthog, complete genome | B263R | 630 | 971 | 71.6% | 4.95e-39 | AY261366 |
| Contig_46493 | 336 | 34.9 | African swine fever virus isolate Malawi Lil-20/1 (1983), complete genome | H359L | 147 | 336 | 77.7% | 3.12e-35 | AY261361 |

**Supplementary Table 3. BLASTn results of SPAdes-assembled contigs containing ASFLI- and other endogenous viral elements detected in the OME/CTVM21 genome**
Shown are hits with the lowest e-value for viral, tick and mobile genetic element-related genes, as obtained by Blastn against the entire NCBI database.

### Supplementary Table 4

**ORFs with length of >500 bp from SPAdes contigs with ASFLI-elements (>500bp)**

| **Contigs** | | | **blastp** | | | | | | | | |
|---|---|---|---|---|---|---|---|---|---|---|---|
| **Designation** | **Length** | **Coverage** | **Protein [Organism]** | **ORF start** | **ORF end** | **Frame** | **AS-length** | **Pairwise identity** | **Positives** | **E-value** | **Accession number** |
| | | | papain [Plasmodium inui San Antonio 1] | 21294 | 20263 | R1 | 344 | 46.0% | 62.0% | 7,00E-09 | XP_008816641.1 |
| | | | RNA-directed DNA polymerase from mobile element jockey, partial [Stegodyphus mimosarum] | 27625 | 28500 | F1 | 292 | 47.0% | 67.0% | 7,00E-65 | KFM61834.1 |
| Contig_885 | 40590 | 7.4 | RNA-directed DNA polymerase from mobile element jockey [Stegodyphus mimosarum] | 29107 | 30402 | F1 | 432 | 39.0% | 60.0% | 2,00E-99 | KFM61834.1 |
| | | | putative RNA-directed DNA polymerase from transposon X-element [Stegodyphus mimosarum] | 30607 | 31185 | F1 | 193 | 50.0% | 67.0% | 2,00E-40 | KFM64962.1 |
| | | | putative RNA-directed DNA polymerase from transposon X-element [Stegodyphus mimosarum] | 32034 | 32942 | F3 | 303 | 32.0% | 50.0% | 4,00E-33 | KFM64962.1 |
| | | | reverse transcriptase [Lasius niger] | 2132 | 1089 | R1 | 348 | 41.0% | 55.0% | 2,00E-38 | KMQ91764.1 |
| | | | reverse transcriptase [Daphnia pulex] | 3481 | 2786 | R2 | 232 | 36.0% | 59.0% | 1,00E-29 | ACB38666.1 |
| | | | Integrase core domain protein [Anaplasma phagocytophilum] | 5476 | 4154 | R2 | 621 | 44.0% | 61.0% | 8,00E-128 | SCV65341.1 |
| Contig_5010 | 16841 | 16.2 | Retrovirus-related Pol polyprotein from transposon opus [Exaiptasia pallida] | 6775 | 5863 | R2 | 301 | 53.0% | 71.0% | 4,00E-103 | KXJ30128.1 |
| | | | PiggyBac transposable element-derived protein 4 [Stegodyphus mimosarum] | 8080 | 8868 | F1 | 263 | 42.0% | 61.0% | 2,00E-45 | KFM62055.1 |
| | | | EP1242L [African swine fever virus Georgia 2007/1] | 10101 | 10847 | F3 | 249 | 87.0% | 90.0% | 3,00E-90 | CBW46719.1 |
| | | | BA71V-EP1242L [African swine fever virus] | 11825 | 12334 | F2 | 170 | 92.0% | 93.0% | 3,00E-76 | AJL34232.1 |
| | | | A859L [African swine fever virus Georgia 2007/1] | 323 | 1111 | F2 | 263 | 69.0% | 77.0% | 4,00E-97 | CBW46706.1 |
| Contig_11845 | 8352 | 4.1 | Pr5-5ELp28 [African swine fever virus] | 2752 | 3354 | F1 | 201 | 53.0% | 70.0% | 8,00E-68 | AAB71363.1 |
| | | | MGF 360-14L [African swine fever virus] | 6267 | 5545 | R1 | 241 | 38.0% | 56.0% | 2,00E-37 | AJL34205.1 |
| Contig_12772 | 7718 | 13.4 | BA71V-H359L (j1L) [African swine fever virus] | 4638 | 4015 | R3 | 308 | 81.0% | 90.0% | 2,00E-114 | AJL34126.1 |
| Contig_17249 | 5489 | 4.2 | BA71V-H359L (j1L) [African swine fever virus] | 5424 | 4534 | R3 | 297 | 78.0% | 88.0% | 3,00E-140 | AJL34126.1 |
| Contig_18126 | 5152 | 26.6 | PREDICTED: piggyBac transposable element-derived protein 3-like isoform X1 [Austrofundulus limnaeus] | 507 | 1 | R2 | 169 | 36.0% | 50.0% | 9,00E-24 | XP_013865069.1 |
| | | | BA71V-C962R [African swine fever virus] | 244 | 897 | F1 | 218 | 76.0% | 86.0% | 9,00E-108 | AJL34250.1 |
| Contig_21116 | 3824 | 4.2 | TPA_inf: pogo transposable element with KRAB domain [Amblyomma variegatum] | 1895 | 2581 | F2 | 229 | 40.0% | 63.0% | 2,00E-32 | DAA34512.1 |
| | | | pogo transposable element with KRAB domain, partial [Acanthochromis polyacanthus] | 2588 | 3148 | F2 | 187 | 55.0% | 70.0% | 2,00E-63 | XP_022078454.1 |
| Contig_22316 | 3339 | 7.3 | piggyBac transposable element-derived protein 4-like [Anoplophora glabripennis] | 1379 | 2650 | F2 | 424 | 46.0% | 62.0% | 1,00E-103 | XP_018562168.1 |
| Contig_24457 | 2653 | 7.9 | BA71V-EP1242L [African swine fever virus] | 872 | 315 | R2 | 186 | 62.0% | 67.0% | 8,00E-39 | AJL34067.1 |
| Contig_35077 | 1763 | 23.9 | BA71V-F778R [African swine fever virus] | 532 | 1083 | F1 | 184 | 88.0% | 91.0% | 3,00E-92 | AJL34059.1 |

**ORFs with length of >500 bp from SPAdes contigs with ASFLI-elements (150-500bp)**

| **Contigs** | | | **blastp** | | | | | | | | |
|---|---|---|---|---|---|---|---|---|---|---|---|
| **Designation** | **Length** | **Coverage** | **Protein [Organism]** | **ORF start** | **ORF end** | **Frame** | **AS-length** | **Pairwise identity** | **Positives** | **E-value** | **Accession number** |
| Contig_1333 | 34743 | 6.0 | oligopeptide transporter, putative [Ixodes scapularis] | 4792 | 5319 | F1 | 176 | 40.0% | 47.0% | 4,00E-18 | XP_002414468.1 |
| Contig_5756 | 15286 | 5.3 | DNA polymerase type B, partial [Nuttalliella namaqua] | 4235 | 4795 | F2 | 187 | 36.0% | 54.0% | 2,00E-11 | AHN53447.1 |
| | | | DNA polymerase type B, partial [Nuttalliella namaqua] | 4692 | 5252 | F3 | 187 | 45.0% | 64.0% | 9,00E-40 | AHN53447.1 |
| Contig_6417 | 14138 | 5.8 | piggyBac transposable element-derived protein 2-like [Astyanax mexicanus] | 1344 | 844 | R3 | 167 | 46.0% | 66.0% | 6,00E-04 | XP_022528824.1 |
| | | | PREDICTED: piggyBac transposable element-derived protein 4 [Tribolium castaneum] | 2593 | 3120 | F1 | 176 | 50.0% | 65.0% | 3,00E-23 | XP_008200478.1 |
| Contig_8979 | 10836 | 16.6 | PREDICTED: piggyBac transposable element-derived protein 4-like [Acyrthosiphon pisum] | 3196 | 3855 | F1 | 220 | 54.0% | 75.0% | 2,00E-65 | XP_008189097.1 |
| | | | hypothetical protein IscW_ISCW020689 [Ixodes scapularis] | 4915 | 5562 | F1 | 216 | 52.0% | 62.0% | 1,00E-19 | XP_002406758.1 |
| Contig_10451 | 9433 | 18.9 | hypothetical protein IscW_ISCW004909 [Ixodes scapularis] | 6476 | 6997 | F2 | 174 | 33.0% | 49.0% | 9,00E-09 | XP_002408708.1 |
| | | | Retrovirus-related Pol polyprotein from transposon 17.6 [Exaiptasia pallida] | 7884 | 8729 | F3 | 282 | 47.0% | 64.0% | 5,00E-48 | KXJ05384.1 |
| | | | Retrovirus-related Pol polyprotein from transposon 17.6, partial [Stegodyphus mimosarum] | 3629 | 4159 | F2 | 177 | 45.0% | 65.0% | 5,00E-40 | KFM82237.1 |
| Contig_11363 | 8706 | 7.2 | Retrovirus-related Pol polyprotein from transposon opus [Exaiptasia pallida] | 4391 | 5842 | F2 | 484 | 38.0% | 58.0% | 5,00E-90 | KXJ30128.1 |
| | | | Retrotransposable element Tf2 protein type 1 [Larimichthys crocea] | 5685 | 7586 | F3 | 634 | 41.0% | 59.0% | 6,00E-136 | KKF31357.1 |
| Contig_12106 | 8169 | 11.0 | NS1 [Lupine feces-associated densovirus 2] | 2912 | 2178 | R2 | 245 | 34.0% | 54.0% | 8,00E-31 | ASM93489.1 |
| Contig_13936 | 7045 | 7.1 | DNA polymerase type B, partial [Nuttalliella namaqua] | 339 | 968 | F3 | 210 | 28.0% | 51.0% | 1,00E-17 | AHN53447.1 |
| | | | DNA polymerase type B, partial [Nuttalliella namaqua] | 2651 | 3175 | F2 | 175 | 49.0% | 70.0% | 6,00E-45 | AHN53447.1 |
| Contig_16776 | 5688 | 5.4 | B263R [African swine fever virus Georgia 2007/1] | 4236 | 5003 | F3 | 256 | 75.0% | 85.0% | 4,00E-72 | CBW46753.1 |
| Contig_18735 | 4920 | 9.0 | gag protein, partial [Nuttalliella namaqua] | 2843 | 3982 | F2 | 380 | 34.0% | 55.0% | 6,00E-34 | AHN53425.1 |
| | | | Transposon Ty3-I Gag-Pol polyprotein [Folsomia candida] | 3790 | 4821 | F1 | 344 | 27.0% | 49.0% | 3,00E-22 | OXA45137.1 |
| Contig_32100 | 2079 | 4.0 | Transposase, ISXO2-like domain-containing protein [Strongyloides ratti] | 935 | 435 | R2 | 167 | 33.0% | 55.0% | 4,00E-10 | CEF70351.1 |

**Supplementary Table 4. BLASTp results of all ORFs >500 bp on SPAdes-assembled contigs containing ASFLI- and other endogenous viral elements detected in the OME/CTVM21 genome**
Shown are hits with the lowest e-value for viral-, tick and mobile genetic element-related proteins, as obtained by Blastp of ORFs >500 bp against the entire NCBI database.

### Supplementary Table 5

| | | | SPAdes contigs with ASFLI-elements from O.porcinus France and Kenya (>150bp) | | | | | | |
|---|---|---|---|---|---|---|---|---|---|
| | Contigs | | | blastn-results | | | | | |
| Designation | Length | Coverage | Organism/Gene | ASFV gene | Query start | Query end | Pairwise identity | E-value | Accession number |
| NODE_104223_OPF | 579 | 1.6 | African swine fever virus Georgia 2007/1 complete genome | EP1242L | 315 | 579 | 76.9% | 1.87e-55 | FR682468 |
| NODE_94830_OPF | 629 | 2.0 | African swine fever virus isolate Kenya 1950, complete genome | F778R | 1 | 433 | 83.2% | 7.10e-131 | AY261360 |
| NODE_38202_OPF | 1556 | 2.7 | African swine fever virus strain Ken05/Tk1, complete genome | G1211R | 1144 | 1552 | 81.2% | 3.76e-101 | KM111294 |
| NODE_11526_OPF | 3089 | 2.8 | African swine fever virus isolate Warmbaths, complete genome | EP1242L | 916 | 2262 | 81.9% | 0,00E+00 | AY261365 |
| NODE_8981_OPF | 3455 | 2.7 | African swine fever virus isolate Warthog, complete genome | B263R | 1993 | 2380 | 70.1% | 7.95e-38 | AY261366 |
| NODE_1447_OPF | 6577 | 3.5 | African swine fever virus isolate Kenya 1950, complete genome | C962R | 4014 | 5083 | 77.8% | 0,00E+00 | AY261360 |
| NODE_1330_OPF | 6757 | 4.4 | African swine fever virus isolate Pretorisuskop/96/4, complete genome | I226R-I243L | 1788 | 2917 | 72.4% | 0,00E+00 | AY261363 |
| NODE_955_OPF | 7421 | 4.6 | African swine fever virus isolate Kenya 1950, complete genome | EP1242L | 3 | 1039 | 79.7% | 0,00E+00 | AY261360 |
| NODE_132770_OPK | 431 | 5.5 | African swine fever virus isolate Mkuzi 1979, complete genome | I226R | 191 | 431 | 71.5% | 1.77e-22 | AY261362 |
| NODE_82440_OPK | 513 | 1.5 | African swine fever virus strain Ken05/Tk1, complete genome | G1211R | 2 | 250 | 79.6% | 4.40e-50 | KM111294 |
| NODE_78000_OPK | 523 | 5.3 | African swine fever virus isolate Kenya 1950, complete genome | C962R | 345 | 503 | 73.0% | 3.0e-8 | AY261360.1 |
| NODE_59972_OPK | 576 | 2.7 | African swine fever virus isolate Warmbaths, complete genome | EP1242L | 204 | 576 | 84.0% | 3.0e-115 | AY261365.1 |
| NODE_28471_OPK | 777 | 2.5 | African swine fever virus Georgia 2007/1 complete genome | IGR | 144 | 447 | 73.0% | 7.0e-48 | FR682468.1 |
| NODE_15700_OPK | 1104 | 2.7 | African swine fever virus isolate Malawi Lil-20/1 (1983), complete genome | C962R | 355 | 976 | 80.0% | 9.0e-163 | AY261361.1 |
| NODE_2992_OPK | 2605 | 2 | African swine fever virus isolate Warmbaths, complete genome | EP1242L | 1016 | 1507 | 84.0% | 3.0e-154 | AY261365.1 |
| NODE_1713_OPK | 3378 | 3.4 | African swine fever virus isolate Pretorisuskop/96/4, complete genome | DP79L-D339L-I226R-I243L | 1393 | 2536 | 71.0% | 2.0e-170 | AY261363.1 |
| NODE_1713_OPK | 3378 | 3.4 | African swine fever virus isolate Pretorisuskop/96/4, complete genome | DP79L-D339L-I226R-I243L | 593 | 1356 | 67.0% | 8.0e-61 | AY261363.1 |
| NODE_1089_OPK | 4100 | 15.1 | African swine fever virus isolate Malawi Lil-20/1 (1983), complete genome | B263R | 477 | 815 | 69.0% | 1.0e-35 | AY261361.1 |
| NODE_769_OPK | 4758 | 5.6 | African swine fever virus isolate 47/Ss/2008, complete genome | EP1242L | 390 | 792 | 77.0% | 8.0e-95 | KX354450.1 |

**Supplementary Table 5. BLASTn results of SPAdes-assembled contigs from O*rnithodoro porcinus* France and Kenya17 containing ASFLI-elements.**
Shown are hits with the lowest e-value for ASFV-genes as obtained by Blastn against the entire NCBI database.

### Supplementary Table 6

| Contigs | | | | Mapping | | | |
|---|---|---|---|---|---|---|---|
| **Designation** | **Length** | **Identified by** | **Mapped against ASFLI-contig** | **ASFLI-gene** | **Hit start** | **Hit end** | **Pairwise identity** |
| AGL001_NODE1 | 7406 | Mapping/Assembly | NODE_5010_length_16841_cov_16.283475 | EP1242L-EP84R-EP424R-F778R-F165R-F1055L_NODE_5010_2 | 1 | 7406 | 99,61% |
| AGL001_NODE2 | 546 | Mapping/Assembly | NODE_35077_length_1763_cov_23.911369 | F778R_NODE_35077 | 1 | 546 | 99,08% |
| AGL001_NODE3 | 670 | Mapping/Assembly | NODE_29175_length_2475_cov_6.846252 | H359L_NODE_29175 | 1 | 670 | 99,69% |
| AGL001_NODE4 | 5185 | Mapping/Assembly | NODE_18126_length_5152_cov_26.682786 | A151R-A104R-A240R-A224L-NODE_18126 | 1 | 5158 | 99,22% |
| AGL001_NODE5 | 3652 | Mapping/Assembly | NODE_17249_length_5489_cov_4.257553 | I267L-G1211R-CP123L-H359L-P1192R-Q706L-Q509L-NODE_17249 | 1 | 3652 | 97,24% |
| AGL001_NODE6 | 2713 | Mapping/Assembly | NODE_12772_length_7718_cov_13.408774 | D1133L-H171R-H24R-NODE_12772 | 1 | 2713 | 98,71% |
| AGL001_NODE7 | 386 | Mapping/Assembly | NODE_28076_length_2653_cov_7.981789 | C147L-C62L-C962R-C475-NODE_28076 | 1/154 | 77/386 | 99,74% |
| AGL001_NODE8 | 274 | Mapping/Assembly | NODE_24457_length_3339_cov_7.382005 | EP1242L-NODE_24457 | 1 | 274 | 100,00% |
| AGL001_NODE9 | 249 | Mapping/Assembly | NODE_24457_length_3339_cov_7.382005 | EP1242L-NODE_24457 | 1 | 249 | 100,00% |
| AGL001_NODE10 | 257 | Mapping/Assembly | NODE_21116_length_4152_cov_12.849193 | C962R-NODE_21116 | 1 | 257 | 98,10% |
| AGL001_NODE11 | 252 | Mapping/Assembly | NODE_19570_length_4637_cov_5.308426 | CP530R-CP80R-CP312R-NODE_19570 | 1 | 252 | 96,80% |
| AGL001_NODE12 | 577 | Mapping/Assembly | NODE_19570_length_4637_cov_5.308426 | CP530R-CP80R-CP312R-O174-NODE_19570 | 1/421 | 213/577 | 100,00% |
| AGL001_NODE13 | 315 | Mapping/Assembly | NODE_19570_length_4637_cov_5.308426 | CP312R-O174-NODE_19570 | 1 | 315 | 97,10% |
| AGL001_NODE14 | 370 | Mapping/Assembly | NODE_19570_length_4637_cov_5.308426 | CP312R-O174-NODE_19570 | 1 | 370 | 98,40% |
| AGL001_NODE15 | 220 | Mapping/Assembly | NODE_15264_length_6370_cov_4.023066 | B175L-B263R-B66L-G1340L-NODE_15264 | 1 | 220 | 100,00% |
| AGL001_NODE16 | 1007 | Mapping/Assembly | NODE_15264_length_6370_cov_4.023066 | B175L-B263R-B66L-G1340L-C962R-NODE_15264 | 1 | 329 | 94,90% |
| AGL001_NODE17 | 334 | Mapping/Assembly | NODE_11845_length_8352_cov_4.102979 | MGF360-15R-NODE_11545 | 1 | 334 | 96,10% |
| AGL001_NODE18 | 233 | Mapping/Assembly | NODE_11845_length_8352_cov_4.102979 | MGF360-15R-NODE_11545 | 1 | 233 | 92,70% |
| MPA001_NODE1 | 3380 | Mapping/Assembly | NODE_955_OPF_length_7421_cov_4.631391 | EP1242L-NODE_955_OPF | 1 | 3380 | 99,17% |
| MPA001_NODE2 | 641 | Mapping/Assembly | NODE_94830_OPF_length_629_cov_2.000000 | MGF360-15R-F778R-NODE_94830_OPF | 1 | 641 | 100,00% |
| MPA001_NODE3 | 1803 | Mapping/Assembly | NODE_8981_OPF_length_3455_cov_2.736478 | B263R-NODE_8981_OPF | 1 | 1803 | 97,95% |
| MPA001_NODE4 | 579 | Mapping/Assembly | NODE_59972_OPK_length_576_cov_2.668151 | EP1242L_NODE_59972_OPK | 1 | 579 | 97,94% |
| MPA001_NODE5 | 446 | Mapping/Assembly | NODE_2992_OPK_length_2605_cov_2.006051 | EP1242L_NODE_2992_OPK | 1 | 446 | 91,03% |
| MPA001_NODE6 | 772 | Mapping/Assembly | NODE_28471_OPK_length_777_cov_2.523077 | IGR_NODE_28471_OPK | 1 | 772 | 94,84% |
| MPA001_NODE7 | 496 | Mapping/Assembly | NODE_1713_OPK_length_3378_cov_3.386343 | DP79L-D339L-NODE_1713_OPK | 1 | 496 | 98,18% |
| MPA001_NODE8 | 781 | Mapping/Assembly | NODE_1713_OPK_length_3378_cov_3.386343 | I226R-I243L-NODE_1713_OPK | 1 | 781 | 99,23% |
| MPA001_NODE9 | 571 | Mapping/Assembly | NODE_15700_OPK_length_1104_cov_2.670420 | C962R-NODE_15700_OPK | 1 | 571 | 92,96% |
| MPA001_NODE10 | 2551 | Mapping/Assembly | NODE_1447_OPF_length_6577_cov_3.541286 | C962R-NODE_1447_OPF | 1 | 2551 | 96,26% |
| MPA001_NODE11 | 3209 | Mapping/Assembly | NODE_1330_OPF_length_6757_cov_4.418703 | I226R-I243L-NODE_1330_OPF | 1 | 3209 | 97,70% |
| MPA001_NODE12 | 355 | Mapping/Assembly | NODE_132770_OPK_length_431_cov_5.470395 | I226R-NODE_132770_OPK | 1 | 355 | 98,31% |
| MPA001_NODE13 | 3100 | Mapping/Assembly | NODE_11526_OPF_length_3089_cov_2.836597 | G1211R-EP1242L-F105L-NODE_11526_OPF | 1 | 3100 | 99,77% |
| MPA001_NODE14 | 265 | Mapping/Assembly | NODE_104223_OPF_length_579_cov_1.612832 | EP1242L-NODE_104223_OPF | 1 | 265 | 100,00% |

**Supplementary Table 6. SPAdes-assembled contigs from deep sequencing data from libraries AGL001 and MPA001, as generated from museum-stored ticks**
Data from libraries from museum-stored tick were mapped against ASFLI-element-containing databases of *Ornithoidoros moubata* (AGL001) and *Ornithodoros porcinus* (MPA001) using Bowtie2 (v.2.3.4.3) with default parameters. Subsequently, mapped reads were assembled using SPAdes and anligned to the corresponding ASFLI-contigs using MAFFT (v. 7.388) in Geneious.

### Supplementary Table 7

| | Sample | ASFV-like sequences | | | | | | | ASFV | PAN-Asfar | Tick control | | | |
|---|---|---|---|---|---|---|---|---|---|---|---|---|---|---|
| | | ASFLIP1192R | | ASFLI-O61R | ASFLI-A104R | ASFLI-A276R | ASFLI-EP1242L | ASFLI-H171R | ASFV B646L | F1055L | *O. moubata* 16S rDNA-gene | *O. moubata* Enolase | | *O.erraticus* Subolesin |
| | | Cq | Cn | Cq | Cq | Cq | Cq | Cq | Cq | Cq | Cq | Cq | Cn | Cq |
| Tick cells | OME/CTVM21 | 22,29 | 147976 | 20,87 | 18,47 | 19,63 | 18,07 | 21,19 | no Cq | 17,84 | 11,60 | 22,27 | 143606 | N/A |
| | OME/CTVM22 | 21,58 | 238530 | 20,36 | 20,26 | 19,52 | 18,67 | 20,61 | no Cq | 19,22 | 11,47 | 21,70 | 211702 | N/A |
| | OME/CTVM24 | 23,77 | 55109 | 22,48 | 22,16 | 21,42 | 20,96 | 23,19 | no Cq | 21,20 | 14,29 | 23,65 | 56266 | N/A |
| | OME/CTVM25 | 28,59 | 2188 | 22,54 | 22,50 | 21,22 | 21,44 | 23,74 | no Cq | 26,19 | 14,90 | 29,12 | 1400 | N/A |
| | OME/CTVM26 | 25,18 | 21390 | 34,20 | 32,29 | 31,64 | 31,67 | 34,72 | no Cq | 22,31 | 23,70 | 26,03 | 11314 | N/A |
| | OME/CTVM27 | 21,30 | 286687 | 20,16 | 17,41 | 18,82 | 17,44 | 20,23 | no Cq | 17,34 | 11,56 | 21,29 | 277649 | N/A |
| Ticks | *O. moubata* Berlin | 23,94 | 49025 | 23,38 | 20,86 | 21,83 | 20,78 | 23,36 | no Cq | 20,61 | 17,03 | 25,33 | 18157 | N/A |
| | *O. moubata* France | 24,44 | 35015 | 24,92 | 20,67 | 20,15 | 19,19 | 23,24 | no Cq | 20,62 | 18,69 | 25,94 | 11975 | N/A |
| | *O. moubata* Pirbright | 28,08 | 2515 | 28,35 | 27,76 | 26,65 | 26,24 | 28,04 | no Cq | 28,48 | 22,01 | 29,96 | 642 | N/A |
| | *O. moubata* Spain | 26,02 | 9447 | 26,20 | 25,77 | 24,10 | 24,82 | 26,16 | no Cq | 26,80 | 18,79 | 28,50 | 1612 | N/A |
| | *O. porcinus* Kenya 2017 | no Cq | 0 | no Cq | no Cq | no Cq | no Cq | no Cq | no Cq | no Cq | 20,18 | 28,76 | 1777 | N/A |
| | *O. porcinus* kenya 2014 | no Cq | 0 | no Cq | no Cq | no Cq | no Cq | no Cq | no Cq | no Cq | 29,77 | no Cq | 0 | N/A |
| | *O. savigny* Nigeria | no Cq | 0 | no Cq | no Cq | no Cq | no Cq | no Cq | no Cq | no Cq | 27,32 | no Cq | 0 | N/A |
| | *O. erraticus* Portugal | no Cq | 0 | no Cq | no Cq | no Cq | no Cq | no Cq | no Cq | no Cq | no Cq | no Cq | 0 | 26,07 |
| ASFV | ASFV Ken06.bus | no Cq | 0 | no Cq | no Cq | no Cq | no Cq | no Cq | 24,67 | 31,82 | N/A | no Cq | 0 | N/A |
| | ASFV ken05/tk1 | N/A | N/A | N/A | N/A | N/A | N/A | N/A | N/A | 24,83 | N/A | N/A | N/A | N/A |
| | ASFV ken.rie1 | N/A | N/A | N/A | N/A | N/A | N/A | N/A | N/A | 21,72 | N/A | N/A | N/A | N/A |
| | ASFV Estonia | N/A | N/A | N/A | N/A | N/A | N/A | N/A | N/A | 31,71 | N/A | N/A | N/A | N/A |
| | ASFV Armenia08 | N/A | N/A | N/A | N/A | N/A | N/A | N/A | N/A | 30,34 | N/A | N/A | N/A | N/A |
| | ASFV Netherlands | N/A | N/A | N/A | N/A | N/A | N/A | N/A | N/A | 28,52 | N/A | N/A | N/A | N/A |
| | ASFV Sardinia | N/A | N/A | N/A | N/A | N/A | N/A | N/A | N/A | 21,02 | N/A | N/A | N/A | N/A |
| | ASFV NHV | N/A | N/A | N/A | N/A | N/A | N/A | N/A | N/A | 14,71 | N/A | N/A | N/A | N/A |

N/A - not tested    Cq - Cycle quantification    Cn - copynumber

**Supplementary Table 7. Results of tick and tick cell line screening for ASFLI-elements by qPCR**

DNA from single ticks and every tick cell line (n=1) was extracted and analysed by qPCR for six ASFV-like genes as described in the 'Material and methods' section. All samples were tested with a qPCR control targeting a tick housekeeping gene to demonstrate successful DNA extraction and presence of tick DNA. Absence of ASFV in samples producing false positive results was proven by an OIE-listed qPCR.

### Supplementary Table 8

| | Sample | ASFV-like sequences | | | | | | | ASFV | Tick control | | | |
|---|---|---|---|---|---|---|---|---|---|---|---|---|---|
| | | ASFLI-P1192R | | ASFLI-O61R | ASFLI-A104R | ASFLI-A276R | ASFLI-EP1242L | ASFLI-H171R | ASFV B646L | *O. moubata* 16S rDNA no RT control | *O. moubata* Enolase | | *O. erraticus* Subolesin |
| | | Cq | Cn | Cq | Cq | Cq | Cq | Cq | Cq | Cq | Cq | Cn | Cq |
| Tick cells | OME/CTVM21 | 25,61 | 17783 | 34,04 | 31,34 | 22,62 | 22,97 | 24,70 | no Cq | 40,35 | 20,02 | 699496 | N/A |
| | OME/CTVM22 | 26,11 | 12804 | 33,08 | 30,80 | 22,41 | 22,77 | 25,08 | no Cq | no Cq | 20,06 | 678583 | N/A |
| | OME/CTVM24 | 27,59 | 4830 | no Cq | 30,89 | 23,05 | 23,41 | 25,87 | no Cq | no Cq | 18,96 | 1393127 | N/A |
| | OME/CTVM25 | 26,32 | 11116 | no Cq | 33,16 | 24,58 | 26,76 | 27,09 | no Cq | no Cq | 20,82 | 412837 | N/A |
| | OME/CTVM26 | 25,76 | 16057 | no Cq | 34,07 | 24,54 | 26,69 | 27,02 | no Cq | no Cq | 21,45 | 273838 | N/A |
| | OME/CTVM27 | 25,46 | 19580 | 34,53 | 31,21 | 24,16 | 25,36 | 24,78 | no Cq | no Cq | 18,86 | 1492178 | N/A |
| Ticks | *O. moubata* Berlin | 23,42 | 75143 | no Cq | 26,59 | 21,24 | 23,28 | 23,91 | no Cq | no Cq | 18,41 | 2003064 | N/A |
| | *O. moubata* France | 24,05 | 49797 | 33,46 | 30,58 | 22,37 | 24,64 | 25,12 | no Cq | no Cq | 19,31 | 1112171 | N/A |
| | *O. porcinus* Kenya 17 | no Cq | 0 | no Cq | no Cq | no Cq | no Cq | no Cq | no Cq | 42,92 | 31,00 | 530 | N/A |
| | *O. porcinus* kenya 14 | no Cq | 0 | no Cq | no Cq | no Cq | no Cq | no Cq | no Cq | no Cq | 34,95 | 50 | N/A |
| | *O. erraticus* Portugal 16 | no Cq | 0 | no Cq | no Cq | no Cq | no Cq | no Cq | no Cq | no Cq | no Cq | 0 | 20,11 |
| ASFV | ASFV Ken06.bus | no Cq | 0 | no Cq | no Cq | no Cq | no Cq | no Cq | 20,86 | N/A | N/A | N/A | N/A |

N/A - not tested     Cq - Cycle quantification     Cn - copynumber

**Supplementary Table 8 Results of tick screening for ASFLI-transcripts by qRT-PCR**

RNA from single *Ornithodoros moubata*, *Ornithodoros porcinus* and *Ornithodoros erraticus* ticks and all six *O. moubata* cell lines (n=1) was extracted and analysed by qRT-PCR as described in the 'Material and methods' section. For all samples, qPCR targeting mitochondrial 16S rDNA was performed without reverse transcriptase checking for residual DNA. Furthermore, all samples were tested with a qRT-PCR control targeting a tick housekeeping gene transcript to demonstrate detectable RNA

### Supplementary Table 9

| | Sample | ASFV titre | 0 dpi NC | 0 dpi | 7 dpi | 14 dpi | 21 dpi | 28 dpi |
|---|---|---|---|---|---|---|---|---|
| ASFV Ken.rie1 (GT X) | *O. moubata* Berlin (1) | | | 36,36 | no Cq | no Cq | no Cq | no Cq |
| | *O. moubata* Berlin (2) | | | 37,23 | no Cq | no Cq | no Cq | no Cq |
| | *O. moubata* Berlin (3) | | no Cq | 36,50 | no Cq | no Cq | no Cq | no Cq |
| | *O. moubata* Berlin (4) | $1 \times 10^4$ HAU/ml | | no Cq | no Cq | no Cq | no Cq | no Cq |
| | *O. moubata* Berlin (5) | | | 37,00 | no Cq | no Cq | no Cq | no Cq |
| | *O. porcinus* (1) | | | no Cq | 34,79 | 25,98 | 27,06 | 26,45 |
| | *O. porcinus* (2) | | no Cq | no Cq | 35,18 | 28,95 | 23,59 | 25,50 |
| | *O. porcinus* (3) | | | 38,66 | 35,54 | 25,07 | 26,52 | 26,31 |
| ASFV Ken.rie1 (GT X) | *O. moubata* Berlin (1) | | | 30.48 | 25.56 | no Cq | no Cq | 35.91 |
| | *O. moubata* Berlin (2) | | no Cq | 30.33 | 37.14 | no Cq | no Cq | 34.45 |
| | *O. moubata* Berlin (3) | $1 \times 10^6$ HAU/ml | | 29.75 | no Cq | 37.22 | 35.24 | 23.38 |
| | *O. porcinus* (1) | | no Cq | 30.33 | 27.39 | 24.65 | 18.78 | 20.66 |
| | *O. porcinus* (2) | | | 29.57 | 20.65 | 18.19 | 17.96 | 19.34 |
| ASFV ken06.bus (GT IX) | *O. moubata* Berlin (1) | | | 36,59 | no Cq | 33,14 | no Cq | no Cq |
| | *O. moubata* Berlin (2) | | | 34,07 | no Cq | 30,40 | no Cq | no Cq |
| | *O. moubata* Berlin (3) | | no Cq | 34,50 | no Cq | no Cq | 34,37 | no Cq |
| | *O. moubata* Berlin (4) | $1 \times 10^5$ HAU/ml | | no Cq | 32,37 | no Cq | no Cq | no Cq |
| | *O. moubata* Berlin (5) | | | no Cq | no Cq | no Cq | no Cq | no Cq |
| | *O. porcinus* (1) | | | 33,42 | no Cq | no Cq | no Cq | no Cq |
| | *O. porcinus* (2) | | no Cq | no Cq | no Cq | no Cq | no Cq | no Cq |
| | *O. porcinus* (3) | | | 33,35 | no Cq | no Cq | no Cq | no Cq |
| ASFV-Sardinia (GTI) | *O. moubata* Berlin (1) | | | no Cq | no Cq | 35,68 | no Cq | no Cq |
| | *O. moubata* Berlin (2) | | | no Cq | no Cq | 36,94 | no Cq | no Cq |
| | *O. moubata* Berlin (3) | | no Cq | no Cq | no Cq | no Cq | no Cq | no Cq |
| | *O. moubata* Berlin (4) | $1 \times 10^4$ HAU/ml | | 37,46 | no Cq | no Cq | no Cq | 35,38 |
| | *O. moubata* Berlin (5) | | | no Cq | 35,90 | no Cq | no Cq | no Cq |
| | *O. porcinus* (1) | | no Cq | 37,68 | 37,30 | no Cq | 35,27 | no Cq |
| | *O. porcinus* (2) | | | 37,82 | no Cq | no Cq | 38,09 | 35,89 |

NC: Negative control

**Supplementary Table 9. qRT-PCR results of *Ornithodoros* ticks experimentally infected with different ASFV-genotype isolates**
Shown are ASFV-P72 transcript-specific Cq-values of third nymphal stage ticks fed with defibrinated pig blood, containing either $1 \times 10^4$ HAU/ml or $1 \times 10^6$ HAU/ml ASFV-ken.rie1 (GT X) (A-D), $1 \times 10^5$ HAU/ml ASFV-Ken06.bus (GT IX) (E-F) or $1 \times 10^4$ HAU/ml ASFV-Sardinia (GT I) (G-H). Due to the limited number of field ticks available and feeding under artificial conditions, fifteen *Ornithodoros porcinus* ticks were collected in each of three experiments and ten in two experiments while for the laboratory-reared *Ornithodoros moubata*, twenty-five individuals were collected in each of three experiments and fifteen in one experiment. All samples were stored at – 80 °C until RNA-extraction and ASFV transcript-specific qRT-PCR analysis as described in the 'Material and methods' section.

### Supplementary Table 10

| ASFV Isolate | Country of origin | *O.porcinus* siRNA (22nt) 22 nt seed unique | raw | piRNA (28-29 nt) 28 nt seed unique | raw |
|---|---|---|---|---|---|
| Total reads | | 595.414 | 6.101.035 | 978.297 | 7.757.885 |
| *O.porcinus* | ASFLI db | 1216 | 5396 | 1465 | 17.132 |
| Warthog | Namibia | 11 | 15 | 9 | 48 |
| Warmbaths | South Africa | 11 | 18 | 8 | 47 |
| Georgia 2007/1 | Georgia | 12 | 20 | 7 | 46 |
| Pretorisuskop/96/4 | South Africa | 11 | 15 | 7 | 46 |
| Tengani 62 | Malawi | 9 | 11 | 7 | 46 |
| Malawi Lil-20/1 | Malawi | 9 | 10 | 3 | 30 |
| 47/Ss/2008 | Italy (Sardinia) | 13 | 24 | 4 | 14 |
| BA71 | Spain | 13 | 24 | 4 | 14 |
| Benin 97/1 | Benin | 13 | 24 | 4 | 14 |
| E75 | Spain | 13 | 24 | 4 | 14 |
| Mkuzi 1979 | South Africa | 10 | 16 | 4 | 14 |
| SPEC_57 | South Africa | 8 | 9 | 4 | 14 |
| RSA_2 | South Africa | 7 | 8 | 4 | 14 |
| LIV_5_40 | South Africa | 6 | 6 | 4 | 14 |
| Kenya 1950 | Kenya | 24 | 73 | 1 | 5 |
| Ken05/Tk1 | Kenya | 18 | 45 | 1 | 5 |
| Ken.rie1 | Kenya | 18 | 52 | 1 | 5 |
| Ken06.Bus | Kenya | 22 | 63 | 1 | 1 |
| R8 | Uganda | 22 | 63 | 1 | 1 |

| ASFV Isolate | Country of origin | *O.moubata* siRNA (22nt) 22 nt seed unique | raw | piRNA (28-29 nt) 28 nt seed unique | raw |
|---|---|---|---|---|---|
| Total reads | | 401.952 | 3.330.492 | 657.981 | 3.841.864 |
| *O.moubata* | ASFLI db | 14.461 | 54.703 | 6.400 | 86.023 |
| Ken06.Bus | Kenya | 489 | 2259 | 91 | 1195 |
| R8 | Uganda | 489 | 2259 | 91 | 1195 |
| Georgia 2007/1 | Georgia | 231 | 871 | 38 | 572 |
| Warthog | Namibia | 225 | 864 | 38 | 572 |
| Pretorisuskop/96/4 | South Africa | 224 | 865 | 38 | 571 |
| Warmbaths | South Africa | 227 | 881 | 35 | 567 |
| Malawi Lil-20/1 | Malawi | 215 | 921 | 36 | 562 |
| Tengani 62 | Malawi | 185 | 694 | 31 | 550 |
| Ken05/Tk1 | Kenya | 522 | 2731 | 52 | 435 |
| Ken.rie1 | Kenya | 460 | 2390 | 51 | 433 |
| Kenya 1950 | Kenya | 470 | 2484 | 50 | 432 |
| 47/Ss/2008 | Italy (Sardini | 180 | 663 | 13 | 17 |
| BA71 | Spain | 180 | 663 | 13 | 17 |
| Benin 97/1 | Benin | 176 | 652 | 13 | 17 |
| E75 | Spain | 180 | 663 | 13 | 17 |
| Mkuzi 1979 | South Africa | 192 | 664 | 13 | 17 |
| SPEC_57 | South Africa | 70 | 248 | 4 | 6 |
| RSA_2 | South Africa | 72 | 247 | 3 | 5 |
| LIV_5_40 | South Africa | 50 | 190 | 3 | 5 |

**Supplementary Table 10. Results of small RNA sequencing and mapping against ASFV and ASFLI-elements**

Small RNA was sequenced from *Ornithodoros porcinus* and *Ornithodoros moubata* nymphal stage ticks. After deduplication using BBMap, 22nt siRNA and 28-29nt piRNA fractions were extracted and mapped against ASFV and ASFVLI-elements using Bowtie2.

### Supplementary Table 11

| qPCR | Target | Forward primer | Reverse primer | Probe | Source |
|---|---|---|---|---|---|
| OM2 | O. moubata 16S rDNA | GGA CAA GAA GAC CCT ATG AAT | CCG GTC TGA ACT CAG ATC A | 5'FAM-ACC TCG ATG TTG GAC TTA GGA TAC CTT-BHQ1-3' | Modified from Bastos et al., 2009 |
| OM5 | O. moubata Enolase | TGA CTG TGA CCA ACC CGA AG | AGG CCT ACG ACA ATG TCT GC | 5'FAM-TGG TGC AAT CTT CAG TCT CTC CGC T-BHQ1-3' | This study |
| NAV-1 | ASFLI-A276R | TGA ACT GCT CTG GCT TTC CA | CAG GGC GTA TGC GTA CTC TT | 5'FAM-CCG GCT TTG GTT GAA ATG CTG CTT T-BHQ1-3' | This study |
| NAV-2 | ASFLI-EP1242L | AAG TAC TCG TCC CGC ATG AC | GGA CTC GGT TAT TGT GTC GC | 5'FAM-AAA CCT TAA GCC CGG CGC CTT AAA-BHQ1-3' | This study |
| NAV-3 | ASFLI-P1192R | CGG CAA AGG AGC TGT TTC AC | GGT CCG TTG AAA CAG GCC TA | 5'FAM-ATT TAC TTT GGC GAG GAG TCG GAG C-BHQ1-3' | This study |
| NAV-4 | ASFLI-H171R | AAA CTA GCT CGC TTT CCG CA | TTT CGT CTG GCC ACG CAT | 5'FAM-TTT GGC TGC TGC GCG ATG AGA AG-BHQ1-3' | This study |
| ASFV2 | ASFV-B646L | TGC TCA TGG TAT CAA TCT TAT CG | CCA CTG GGT TGG TAT TCC TC | 5'FAM-TTC CAT CAA AGT TCT GCA GCT CTT-Tamra-3' | Tignon et al., 2011 |
| AL A104R | ASFLI-A104R | AGC CAA CAA TTA CTA AGC AAG AGC | GGC AGG CTT GGC TTT AAC AA | 5'FAM-CGC CCG GAA TTA AAT TCT CAG TGG C-BHQ1-3' | This study |
| AL P12 | ASFLI-P12 | ATG GCA CCT GGT GAT GGT TT | GGA ACA CGT TTC ATT GCT ATT GC | 5'FAM-TGG TAC ATT CCT CGT CAA AAT TCG CA-BHQ1-3' | This study |
| OE3 | O. erraticus-Subolesin | CCT CAG AAC AGC AGC ACA GA | TGG CAT TCA CGC TCC TTC AT | 5'FAM-CCC TGT TCA CCT TTC GCC AAG TAG GG-BHQ1-3' | This study |
| PAN-Asfi | ASFV/ASFLI-F1055L | AAT SGT YTC RGC CGA GGC AAT | TTT AAC CAG RTY GCC GAC AC | 5'FAM-GGC GCY AAG GTG TTT ATA CTR ACG GG-BHQ1-3' | This study |

**Supplementary Table 11. Primer and probe sequences used in this study**

### Supplementary Appendix 1 Phylogenetic analysis and molecular clock analysis

#### Sequence fragments

ASFV-like sequences of the gene EP1242L were found in the samples from four *Ornithodoros* spp. ticks originating from Angola, Tanzania (previously known as German East Africa), Kenya and the *Ornithodoros moubata* cell line OME/CTVM21 (grouped hereafter as "tick samples"), along with fragments of other genes (different fragments in different samples). The entire EP1242L viral gene was not found; however fragments of over 1500bp in length were recovered from each of the samples. EP1242L codes for the DNA-directed RNA polymerase subunit beta in ASFV.

The ASFV-like fragments were aligned to the EP1242L of a reference ASFV genome - ASFV|KM111295|Kenya|Ken06/Bus|2006, and the first 1751 nucleotides in the alignment were used (Table 1). The sequences contained a number of deletions and one insertion as compared to Ken06/Bus.

| **Sample number** | **Sample Name or reference virus** | **Fragment Length** | **Length used** |
|---|---|---|---|
| *(reference)* | *ASFV|KM111295|Kenya|Ken06/Bus|2006* | *3729* | *1751* |
| 1 | Omoubata|Original_Contig|2017 | 2562 | 1641 |
| 2 | AGL001|Angola|1900 | 2561 | 1641 |
| 3 | OME21|tickcells|2018 | 2562 | 1641 |
| 4 | MPA001A|German-East-Africa|1906 | 1630 | 1630 |
| 5 | Oporcinus|Oporc955_FLI|2019 (Kenya) | 1638 | 1638 |

Table 1: Summary of EP1242L fragments derived from ticks and the tick cell line OME/CTVM21 (OME21) used in the analysis

| **Start** | **End** | **Length** | **Ref** | **1** | **2** | **3** | **4** | **5** | **Include** |
|---|---|---|---|---|---|---|---|---|---|
| 9 | 15 | 7 | NN | NN | NN | NN | D1 | D1 | Yes |
| 255 | 270 | 16 | NN | D1 | D1 | D1 | NN | NN | Yes |
| 407 | 412 | 6 | NN | D1 | D1 | D1 | NN | NN | Yes |
| 484 | 484 | 1 | NN | NN | NN | NN | D1 | D1 | Yes |
| 524 | 532 | 9 | NN | D1 | D1 | D1 | D1 | D1 | |
| 563 | 581 | 19 | NN | D1 | D1 | D1 | D1 | D1 | |
| 633 | 648 | 16 | NN | D2 | D2 | D2 | D1 | D1 | Yes |
| 668 | 675 | 8 | NN | D1 | D1 | D1 | D1 | D1 | |
| 742 | 750 | 9 | NN | NN | NN | NN | D1 | D1 | Yes |
| 818 | 822 | 5 | NN | NN | NN | NN | D1 | D1 | Yes |
| 961 | 969 | 9 | NN | NN | NN | NN | D1 | D1 | Yes |
| 1013 | 1023 | 11 | NN | NN | NN | NN | D1 | D1 | Yes |
| 1096 | 1105 | 10 | NN | D1 | D1 | D1 | D1 | D1 | |
| 1385 | 1393 | 9 | NN | D1 | D1 | D1 | D1 | D1 | |
| 1440 | i | | NN | II | II | II | NN | NN | Yes |
| 1456 | 1457 | 2 | NN | NN | NN | NN | D1 | D1 | Yes |
| 1677 | 1683 | 7 | NN | D1 | D1 | D1 | NN | NN | Yes |
| 1730 | 1737 | 8 | NN | D1 | D1 | D1 | D1 | NN | Yes |
| 1747 | 1747 | 1 | NN | NN | NN | NN | D1 | NN | Yes |
| 1751 | 1751 | 1 | NN | NN | NN | NN | NN | D1 | Yes |

Table 2: Indels in the tick sample sequences relative to the start of EP1242L in ASFV|KM111295|Kenya|Ken06/Bus|2006. NN = nucleotide sequence, D1 = deletion region 1, D2 = deletion region 2, II = insertion region 2

### Maximum Likelihood Trees and Relation to ASFV

#### Phylogeny from nucleotide sequences

A maximum likelihood tree of the aligned nucleotide sequences of the samples, together with 17 publicly available ASFV EP1242L sequences (non-integrated) was created using MEGA7 with the Tamura-Nei model, gamma-distributed site-to-site rate variation (four categories), and 100 bootstraps.

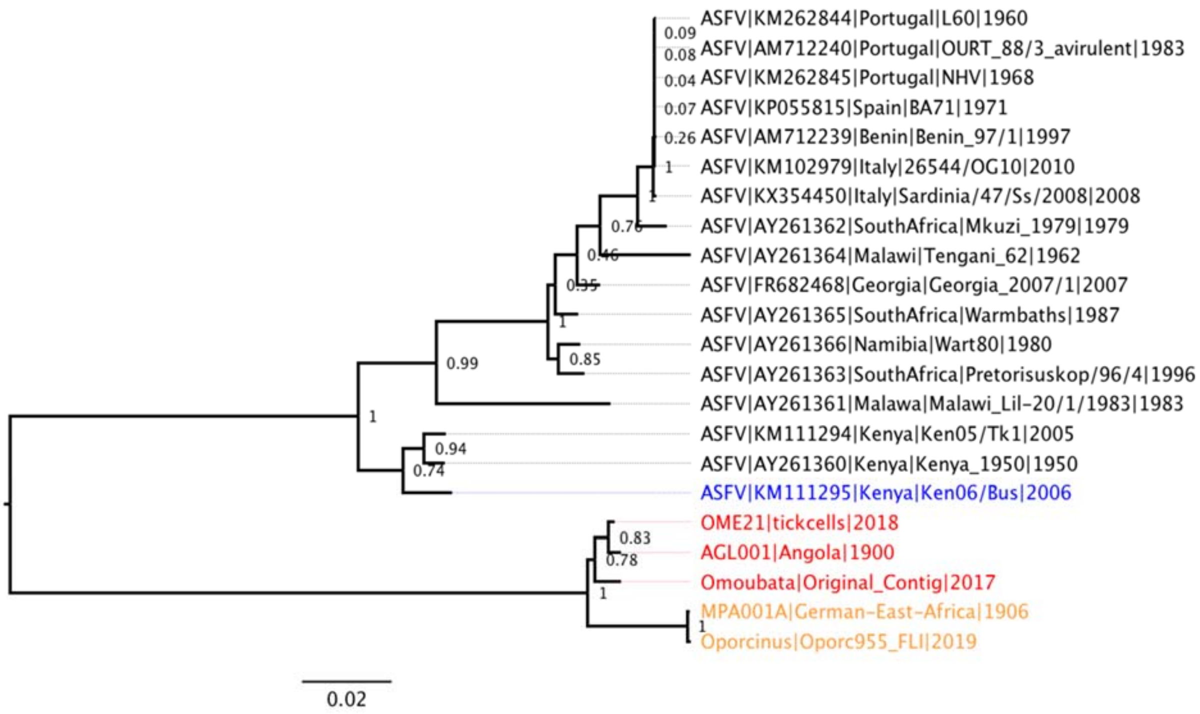

Figure 1: Maximum likelihood tree of the ASFV-like nucleotide fragments from the tick samples and of partial ASFV EP1242L genes (non-integrated)

The tree in Figure 1 shows that the viral gene fragments in the samples from ticks are related to ASFV but appear as outgroups. This is to be expected since the viral gene fragments were integrated into the tick genomes and evolved with the host species, whereas the sequences from free ASFV have been evolving independently in the viral population.

#### Phylogeny from amino acid sequences

To further show the relationship between the ASFV-like sequence fragments and other viruses, a maximum likelihood tree including EP1242L-equivalent sequences from other, more distantly related Megavirales (Pacmanvirus, Faustovirus and Kaumeobavirus) was inferred in MEGA7 from amino acid sequences using the JTT (Jones-Taylor-Thornton) model, gamma-distributed site-to-site rate variation in four categories and 100 bootstraps.

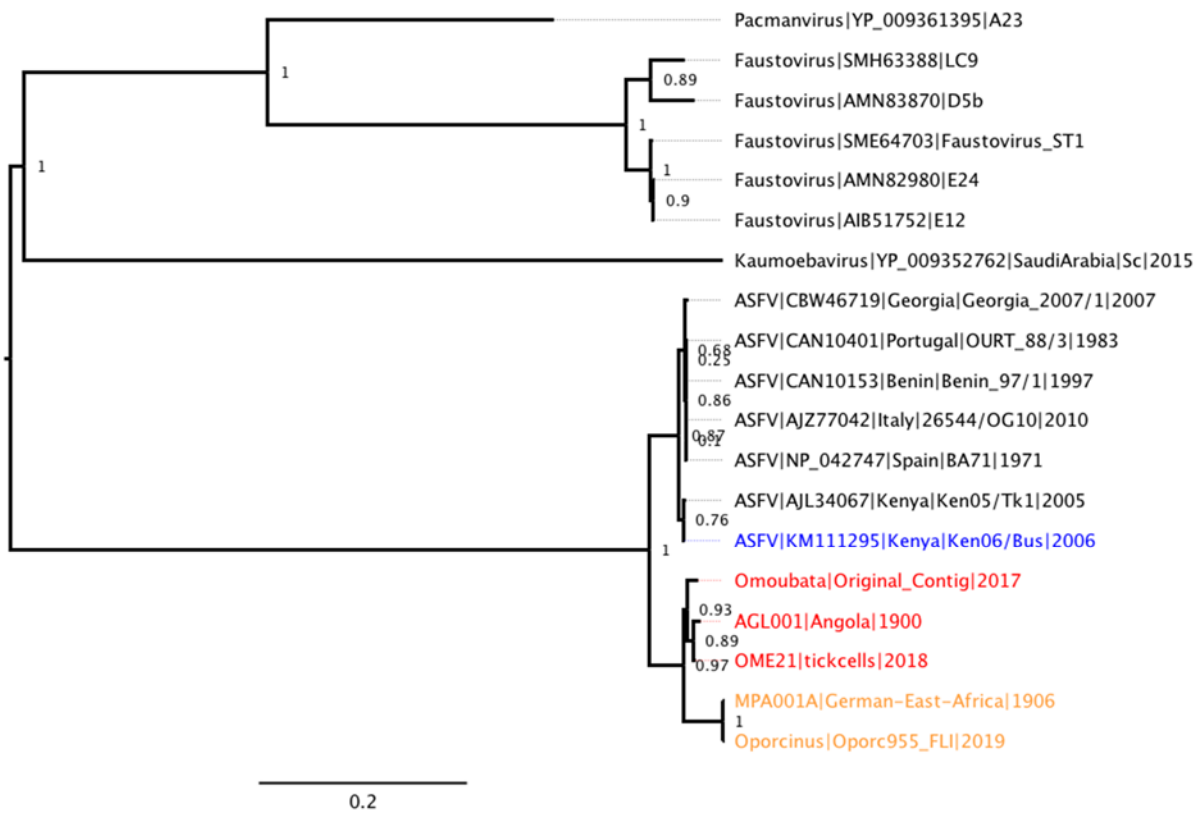

Figure 2: Maximum likelihood tree of the ASFV-like amino acid sequences from the tick samples, and of partial ASFV EP1242L sequences of (non-integrated), Faustovirus, Pacmanvirus and Kaumeobavirus.

### Time-scale estimation

#### Initial estimation

To estimate the time-scale for divergence of the tick samples and ASFV (non-integrated), time-scaled phylogenies were inferred using BEAST 1.10.4 with the following model settings: a Tamura-Nei 93 (TN93) nucleotide substitution model with gamma-distributed site-to-site rate variation in four categories, a strict or uncorrelated relaxed lognormal clock and a constant population size. Initially, the EP1252L sequence fragments from the five ticks together with the Ken06/Bus sequence reference were used. Several priors for the overall clock rate were compared, drawn from a normal or log-normal distribution with means and standard deviations of between 1e-8 substitutions per site and year to 5e-6 per site and year. The set of values for the clock rate priors were chosen to reflect typical DNA molecular clock rates (lower rates for host, higher rates for ASFV).

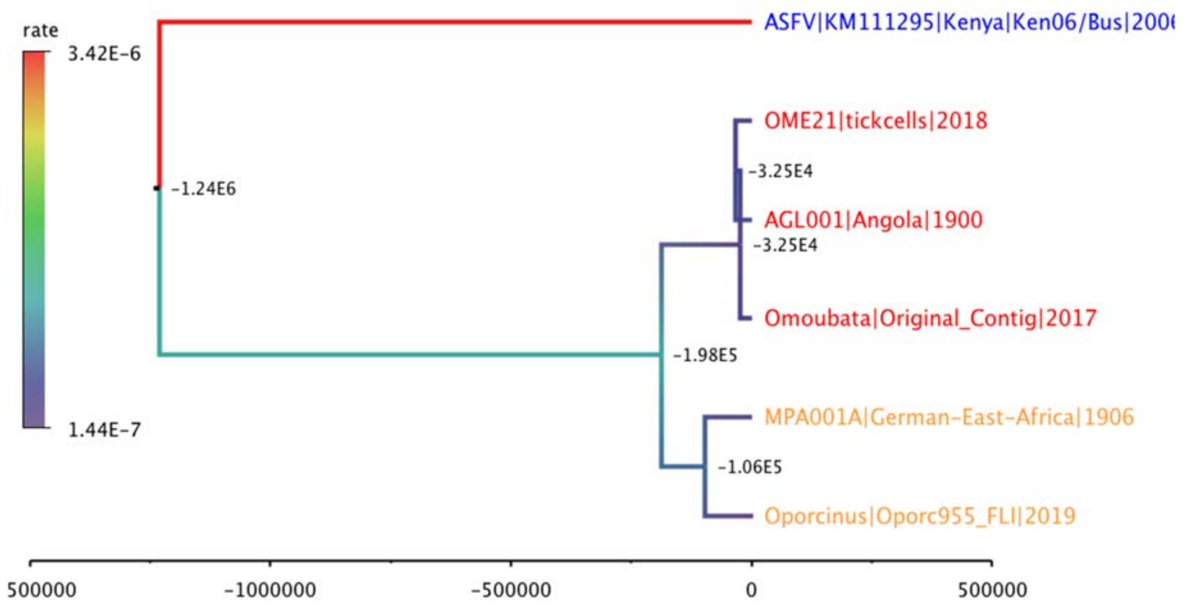

Figure 3: Time-scaled tree for partial EP1242L sequences from tick samples and reference ASFV (non-integrated), inferred using an uncorrelated relaxed log-normal molecular clock and log-normal clock rate prior with mean and standard deviation of 5e-6 substitutions per site and year. Branches are coloured according to the inferred molecular clock rate, dark blue = 1.4e-7 (slow) to red = 3.4e-6 (fast) substitutions/site x year. Node ages are indicated in years BCE.

| | | Root Height | Root Age statistics | | | | | Mean Rate |
|---|---|---|---|---|---|---|---|---|
| Clock | Prior | mean | mean | median | 95% Lower | 95% Upper | ESS | mean |
| Strict | 1e-6 fix | 9.75E+04 | -9.55E+04 | -9.52E+04 | -8.45E+04 | -1.07E+05 | 8389 | 1.00E-06 |
| Strict | 1e-7 fix | 9.76E+05 | -9.74E+05 | -9.71E+05 | -8.69E+05 | -1.09E+06 | 8203 | 1.00E-07 |
| Strict | 1e-8 fix | 9.74E+06 | -9.74E+06 | -9.71E+06 | -8.66E+06 | -1.09E+07 | 8397 | 1.00E-08 |

Table 3: Estimated root height and overall mean clock rate for strict clocks with fixed priors

| | | Root Height | Root Age statistics | | | | | Mean Rate |
|---|---|---|---|---|---|---|---|---|
| Clock | Prior | mean | mean | median | 95% Lower | 95% Upper | ESS | mean |
| RelaxLn | 5e-6 fix | 3.02E+06 | -3.01E+06 | -9.10E+05 | -4.04E+03 | -1.17E+07 | 71 | 7.86E-07 |
| RelaxLn | 1e-6 fix | 9.17E+07 | -9.17E+07 | -1.98E+07 | -2.82E+04 | -3.88E+08 | 112 | 4.94E-08 |
| RelaxLn | 5e-7 fix | 1.07E+09 | -1.07E+09 | -1.64E+08 | -5.91E+04 | -5.16E+09 | 22 | 3.49E-08 |
| RelaxLn | 1e-7 fix | 3.54E+09 | -3.54E+09 | -6.48E+08 | -4.11E+05 | -1.46E+10 | 38 | 3.12E-09 |
| RelaxLn | 5e-8 fix | 2.02E+09 | -2.02E+09 | -1.47E+09 | -4.57E+06 | -5.24E+09 | 100 | 2.48E-10 |
| RelaxLn | 1e-8 fix | 7.79E+12 | -7.79E+12 | -2.16E+12 | -7.30E+08 | -3.51E+13 | 60 | 7.07E-13 |

Table 4: Estimated root height and overall (averaged over all branches) mean clock rate for relaxed clocks with fixed priors of the rate of the relaxed clock (some variation)

| | | Root Height | Root Age statistics | | | | | Mean Rate |
|---|---|---|---|---|---|---|---|---|
| Clock | Prior | mean | mean | median | 95% Lower | 95% Upper | ESS | mean |
| RelaxLn | 5e-6 log | 7.94E+06 | -7.94E+06 | -1.23E+06 | -3.34E+07 | 5.23E+02 | 107 | 7.76E-07 |
| RelaxLn | 5e-6 nor | 2.32E+07 | -2.32E+07 | -3.43E+05 | -6.01E+07 | -1.92E+02 | 80 | 1.18E-06 |
| RelaxLn | 5e-7 log | 1.77E+09 | -1.77E+09 | -8.35E+07 | -8.32E+09 | -5.39E+03 | 86 | 3.55E-08 |
| RelaxLn | 5e-7 nor | 1.40E+09 | -1.40E+09 | -3.58E+07 | -1.45E+09 | -1.90E+04 | 275 | 7.00E-08 |
| RelaxLn | 5e-8 log | 3.86E+11 | -3.86E+11 | -1.26E+10 | -1.71E+12 | -1.50E+06 | 77 | 2.37E-10 |
| RelaxLn | 5e-8 nor | 3.65E+09 | -3.65E+09 | -6.80E+08 | -1.52E+10 | -1.11E+06 | 207 | 7.75E-10 |

Table 5: Estimated root height and overall (averaged over all branches) mean clock rate for relaxed clocks with normal or log-normal priors on the rate of the relaxed clock (most variation)

Estimation using tick samples only

Tables 3-5 and Figure 3 shows that considerable variation in the time-scale estimates is possible, depending on the model settings, especially the choice of clock rate priors. This is partly due to attempting to infer time-scales between integrated and non-integrated ASFV. Furthermore, the effective sample size (ESS) of the MCMC traces from these models were poor (despite running the chain for at least 10,000 steps which would normally be more than sufficient for six taxa), indicating poor model fit.

Therefore, we also inferred time-scaled trees using the five tick samples only. Here we expected that using a strict clock (no variation between branches) and a clock rate prior commensurate with the expected substitution rate for the host DNA would be most appropriate. The aligned nucleotide sequences were used to infer time-scaled trees (excluding the Ken06/Bus reference sequence), using a TN93 model with invariant sites and gamma-distributed rates for the variable sites (four categories), strict clock, constant population size, together with the 15 indel sites which differed between sequences 1-5 (final column of Table 2).

| | | **Root Height** | **Root Age statistics** | | | | | **Mean Rate** |
|---|---|---|---|---|---|---|---|---|
| **Clock** | **Prior** | **mean** | **mean** | **median** | **95% Lower** | **95% Upper** | **ESS** | **mean** |
| Strict | 5e-6 log | 1.11E+04 | -9.13E+03 | -6.04E+03 | -2.73E+04 | 7.49E+02 | 298 | 2.46E-06 |
| Strict | 1e-6 log | 4.50E+04 | -4.30E+04 | -3.01E+04 | -1.25E+05 | -9.44E+02 | 378 | 6.85E-07 |
| Strict | 5e-7 log | 8.08E+04 | -7.88E+04 | -5.77E+04 | -2.18E+05 | -4.10E+03 | 377 | 3.55E-07 |
| Strict | 1e-7 log | 4.50E+05 | -4.48E+05 | -3.20E+05 | -1.25E+06 | -1.68E+04 | 364 | 7.02E-08 |
| Strict | 5e-8 log | 8.91E+05 | -8.89E+05 | -6.24E+05 | -2.48E+06 | -6.66E+04 | 377 | 3.36E-08 |
| Strict | 1e-8 log | 4.70E+06 | -4.70E+06 | -3.22E+06 | -1.31E+07 | -1.42E+05 | 391 | 7.00E-09 |

Table 6: Estimated root height and overall (averaged over all branches) mean clock rate for strict clocks with log-normal priors on the rate of the strict clock for the five ASFV-like sequences from tick samples only.

The difference in fit between these models and priors was assessed by estimating the marginal likelihood using path sampling and stepping stone sampling in BEAST. The results indicate that the log-normal clock rate prior with mean 5e-7 is best, however, the difference in log likelihoods is not very significant between the different prior settings, especially settings 1e-7, 5e-8 and 1e-8 (table 7).

| **Clock** | **Prior** | **Path** | **Stepping** | **Diff Max-Path** | **Diff Max-Stepping** |
|---|---|---|---|---|---|
| Strict | 5e-6 log | -2797.86 | -2797.90 | 0.93 | 0.95 |
| Strict | 1e-6 log | -2798.40 | -2798.53 | 1.47 | 1.58 |
| Strict | 5e-7 log | -2796.93 | -2796.95 | 0.00 | 0.00 |
| Strict | 1e-7 log | -2797.23 | -2797.31 | 0.30 | 0.36 |
| Strict | 5e-8 log | -2797.44 | -2797.51 | 0.51 | 0.56 |
| Strict | 1e-8 log | -2797.81 | -2797.91 | 0.88 | 0.97 |

Table 7: Marginal likelihood estimation using Path Sampling and Stepping Stone Sampling, showing that the log-normal clock rate prior with mean = 5e-7 is best, but not significantly better than the other clock rates.

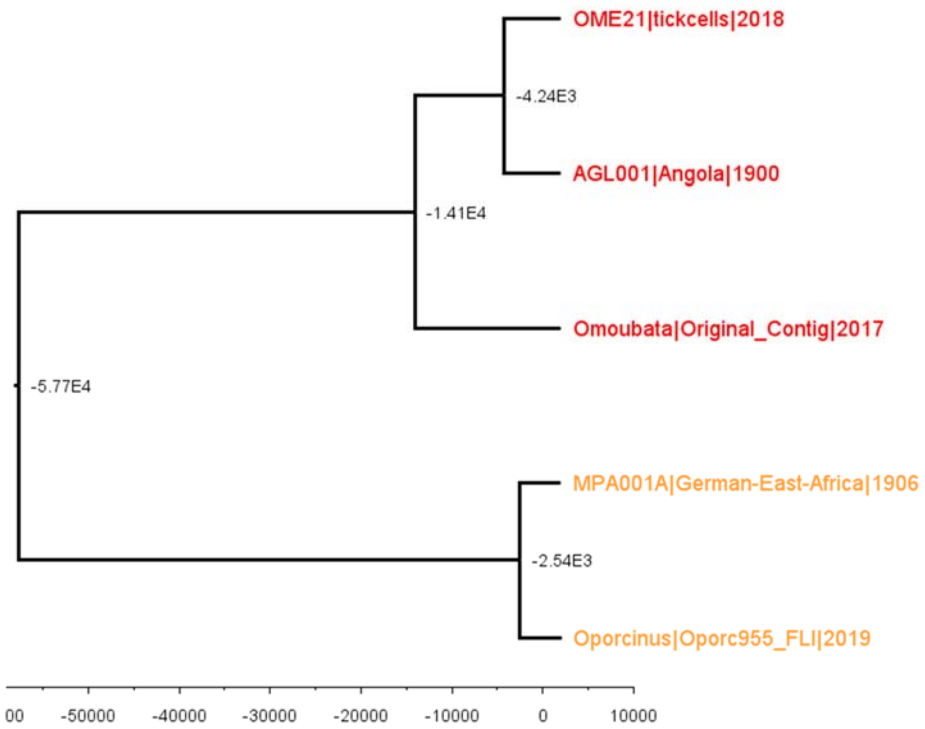

Figure 4: Time-scaled tree for partial EP1242L sequences from tick samples were inferred using a strict molecular clock and log-normal clock rate prior with mean and standard deviation of 5e-7 substitutions per site per year. Node ages are indicated in years BCE.

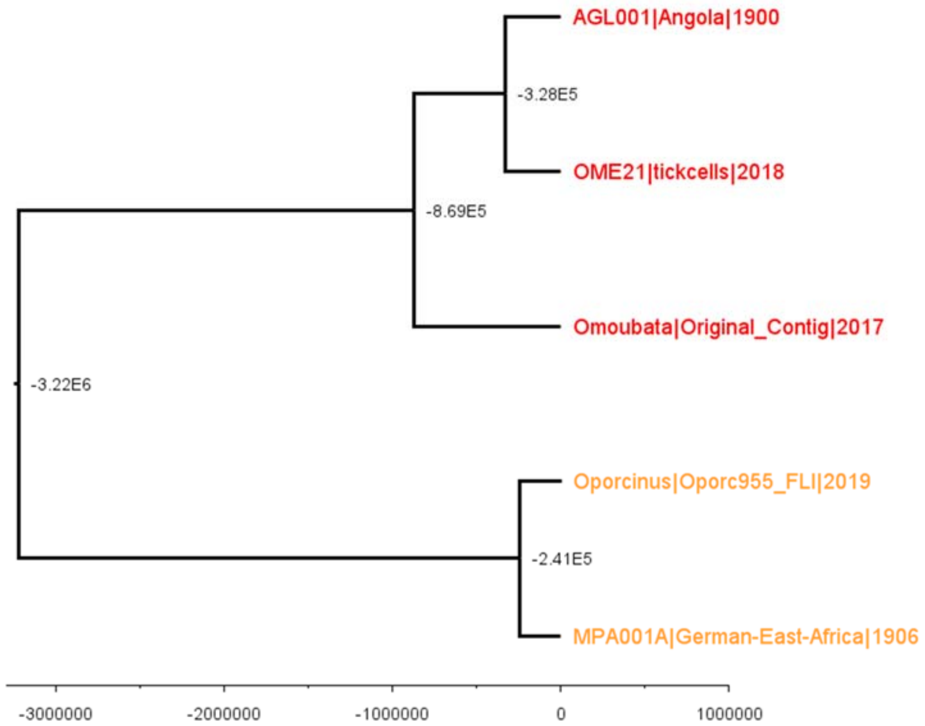

Figure 5: Time-scaled tree for partial EP1242L sequences from tick samples were inferred using a strict molecular clock and log-normal clock rate prior with mean and standard deviation of 1e-8 substitutions per site and year. Node ages are indicated in years BCE.
